# Integrative multi-omics analysis of dietary fibre-induced modulations in the composition and function of chicken caecal microbiota

**DOI:** 10.1101/2025.08.27.672358

**Authors:** Anum Ali Ahmad, Kellie Watson, Farina Khattak, Dominic Kurian, Rachel Kline, Sebastien Guizard, Laura Glendinning

## Abstract

**Background:** The sustainability of poultry farming faces significant challenges due to rising feed costs and competition with human food sources. Dietary fibre offers a promising, cost-effective alternative due to its beneficial impact on gut health. We utilised a multi-omics approach to understand the influence of soluble inulin and insoluble cellulose dietary fibres on the composition and function of caecal microbiota in broilers.

**Results:** High inulin significantly altered caecal microbial composition and promoted broader microbial metabolic adaptations, while high cellulose had a minimal impact.

**Conclusions:** These findings will enhance our understanding of how various fibre types and quantities shape gut microbiota, ultimately leading to improved poultry performance.

## Background

Chickens (Gallus gallus) are the most consumed meat source worldwide, providing high-quality protein, essential amino acids, and micronutrients (Govoni et al. 2021). The global poultry sector is expected to continue to expand as demand for meat is increasing due to growing populations, urbanization, and increasing incomes (Farris et al. 2024). The outlook report of the FAO and OECD predicted a 30% increase in poultry meat production by 2033 (OECD/FAO 2024). This rising demand is causing significant challenges to the poultry sector and putting unprecedented pressure on poultry production systems.

The global human population is expected to exceed 9.7 billion by 2050, demanding a more sustainable food system that uses fewer natural resources with maximum production output (FAO 2021). The primary challenges for sustainable poultry production are feed cost and its availability. High-energy and low-fibre diets, typically based on cereal grains such as maize, soy, and corn, are required by fast-growing genotypes. These diets are not only costly but also directly compete with the human food supply, resulting in feed-food competition (OECD/FAO 2018). Thus, developing alternative and affordable feeding strategies is crucial for food security and environmental sustainability in poultry production (FAO 2023).

A potential solution toward more sustainable poultry production lies in the use of alternative feed resources, including crops or agricultural by-products, that are more affordable and environmentally friendly (Bist et al. 2024). However, these feeds tend to be higher in dietary fibre, which presents a significant challenge for modern broiler chickens. Recently, dietary fibre (NSP-non-starch polysaccharide) has become a major portion of chicken rations, fulfilling roughly 3-4% of the nutritional demands of the birds (Nguyen et al. 2022). NSPs were initially thought to be anti-nutritive (Pérez-Jiménez 2024), but recent studies have shown that soluble and insoluble components of NSP have distinct effects on poultry production performance (Nguyen et al. 2021). Insoluble non-starch polysaccharides (NSP), including cellulose, are known to increase nutrient retention time and enhance feed efficiency, while soluble components, including inulin, pectin, and glucan are known to increase digesta viscosity and reduce nutrient absorption (Marc et al. 2024; Nguyen et al. 2022). Just like ruminants, chickens have a limited intrinsic ability to degrade fibre due to their short digestive tracts and lack of endogenous cellulolytic enzymes. Therefore, they are dependent on their gut microbiota to extract energy from NSPs present in fibrous feed ingredients.

Chicks raised in modern poultry settings lack maternal contact, which is crucial in shaping early gut microbiota (Shterzer et al. 2023). As a result, the gut microbial community of commercially raised birds often lacks key microbial taxa involved in fibre fermentation (Novoa Rama et al. 2023). This decreases the fermentation and energy extraction from a high-fibre diet, as well as their growth performance and feed efficiency. Targeted manipulation of the gut microbiota is an important strategy that can be performed to improve fibre fermentation in poultry.

Chicken caeca harbour the most complex microbial community in the chicken gut, making them the main site for fermenting NSP into volatile fatty acids (VFAs), which serve as an essential energy source for the host (Jamroz et al. 2002; Ocejo et al. 2019). Studies have highlighted both the beneficial and adverse roles of dietary fibre in shaping gut microbiota and its potential impact on bird health through microbial manipulation (Xia et al. 2021; Mirzaie et al. 2012; de Sousa et al. 2025).

To date, most studies have utilised 16S rRNA sequencing approach to explore the impact of NSP on the composition of chicken caecal microbiota (Qiu et al. 2022; de Sousa et al. 2025; Hou et al. 2020). Studies employing shotgun metagenomics to gain deeper insights into microbial composition and functions are scarce and do not capture real-time gene expression or enzymatic activity (Jian et al. 2025). As a result, it offers only a limited understanding of the metabolic processes underlying fibre fermentation. Therefore, integrating multi-omics approaches is essential to unravel the mechanisms by which dietary fibres influence the caecal microbiota and to identify key microbial players and enzymes involved in fibre fermentation. This study is the first to employ a multi-omics approach that includes metagenomics, metatranscriptomics, and metaproteomics to explore the effects of soluble (inulin) and insoluble (cellulose) dietary fibre on the caecal microbial composition and their functional profile in chickens. This study moves beyond descriptions of changes in the chicken gut microbiota upon fibre administration, allowing us to identify the key mechanistic drivers of chicken caecal fibre fermentation.

## Results

### Metagenomic and metatranscriptomic data show differences in dominant microbial genera

Metagenomic (MG) and metatranscriptomic (MT) reads were classified at different taxonomic levels using Kraken2 to characterise the caecal microbial community (Fig. 1, Fig. S1). This approach offers a comprehensive overview of microbial composition and their functional activity, enabling the identification of microbial taxa beyond bacteria and archaea, and providing insight into transcriptionally active members of the microbial community. Most reads were classified as bacteria (MG=86.5% ± 0.67, MT=89.3% ± 0.72), with some belonging to Archaea (MG=0.21% ± 0.08, MT=0.35% ± 0.14) and Eukaryota (MG=0.42% ± 0.03, MT=0.27% ± 0.03) (Fig. S1, values represent mean % ± SEM). Also, 13.4% ± 0.71 of the MG and 10.3% ± 0.70 of the MT reads were not assigned any taxonomy. Bacillota (MG=65.4% ± 2.27, MT=75.2% ± 1.83) and Bacteroidota (MG=14.5% ± 1.84, MT=10.2% ± 1.58) were the most classified phyla, while Faecalibacterium (MG=11.7% ± 1.70, MT=29.3% ± 3.65) and Barnesiella (MG=8.05% ± 1.71, MT=6.16% ± 1.49) were the most classified genera (Fig. S1). We identified Bifidobacterium (2.33% ± 1.16) among the top 10 genera in the MG data, while it was replaced by Agathobaculum (1.85% ± 0.29) in the MT data (Fig. 1). This might be because Bifidobacterium was abundant but not very functionally active.

**Fig. 1.**
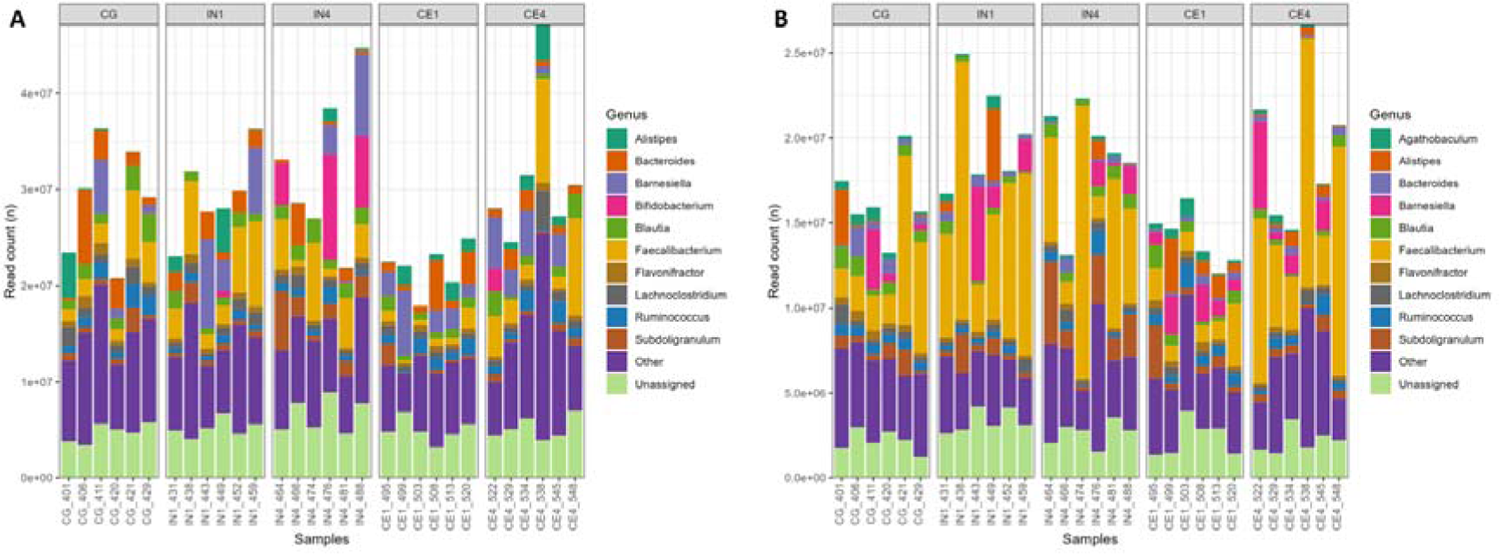
Metagenomics (A) and metatranscriptomics (B) reads classified at the genus level using Kraken2 across different dietary groups. Only the top 10 most abundant taxa are shown, while the remaining taxa are grouped as other. Reads that couldn’t be assigned any taxonomy are classified as unassigned. CG, Control group; IN1, 1% inulin; IN4, 4% inulin; CE1, 1% ARBOCEL; CE4, 4% ARBOCEL

### 514 high-quality MAGs constructed from shotgun metagenome sequencing data

We used metagenomic data to construct metagenome assembled genomes (MAGs), which allows the identification of unculturable, novel microbial species in the caeca. Shotgun metagenomic sequencing of caecal contents samples generated approximately 96.83 ± 3.03 million reads with an average length of 150 bp per sample. Clean reads (85.74 ± 3.34 million mean reads per sample) obtained after quality control and removal of host reads were assembled into 2,991,842 contigs with an average N50 value of 6699 bp. The contigs were binned into MAGs and dereplicated at 95% average nucleotide identity (ANI), a commonly used species-level threshold, to remove redundancy. The MAGs were then filtered to retain only those with contamination ≤5% and completeness ≥80%, resulting in the recovery of 514 high-quality MAGs. The details of each MAG are presented in Table S4. Taxonomic annotation of MAGs revealed that 511 belonged to bacteria while 3 belonged to archaea (Fig. 2).

**Fig. 2.**
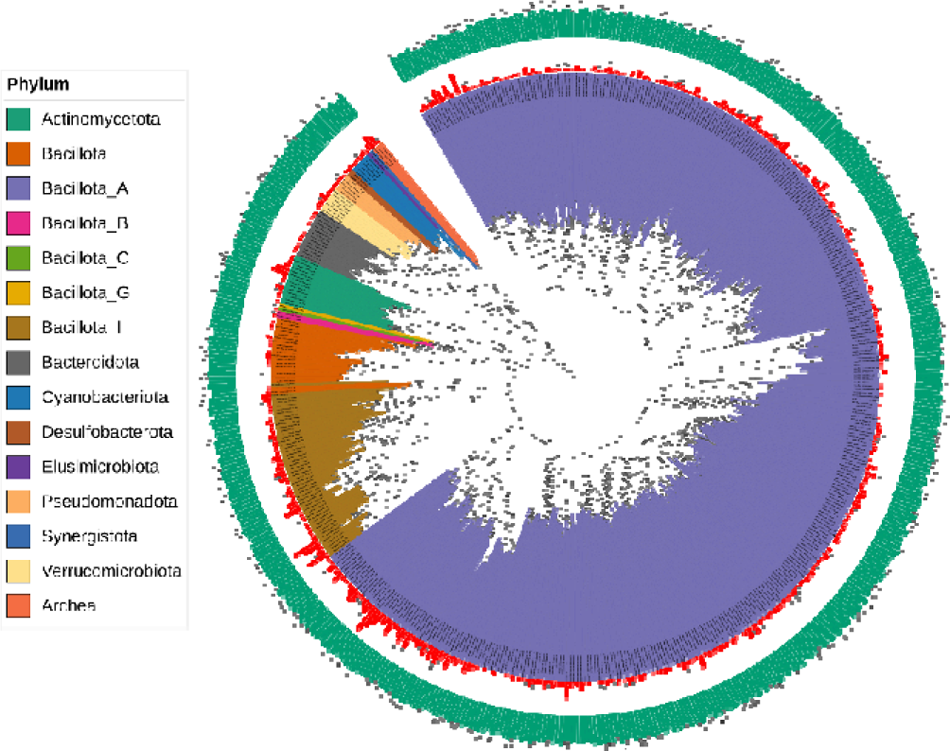
Microbial phylogenetic tree representing 514 metagenome-assembled genomes (MAGs) recovered from the caeca of chickens across all dietary groups. Background colours indicate the phylum to which MAGs belong. The red outer circle indicates contamination levels, while the green outermost circle represents completeness of MAGs.

### High inulin quantity altered the caecal microbial diversity in chickens

We analysed caecal microbial diversity to assess the variety and abundance of microbial taxa within a sample (alpha diversity) and between different samples in response to different dietary fibres. Alpha diversity analysis showed a significant effect of dietary fibre on the caecal microbial community of chickens. Microbial diversity measured by Shannon (Kruskal-Wallis, p= 0.01) and ACE indices (Kruskal-Wallis, p=0.001) showed a decrease in the number of species and their relative abundance in the IN4 group compared to other dietary groups (Fig. 3A and B). Pairwise comparisons revealed significant differences in the IN4 group compared to the CG (p = 0.01) and CE1 (p = 0.01) groups using the Shannon index. Additionally, significant differences in the caecal microbial community of the IN4 group compared to the CE1 (p < 0.001) and CE4 (p < 0.01) groups were observed using the ACE index. Beta diversity analysis based on Bray-Curtis distance showed a significant difference among dietary groups (Adonis2, p < 0.001). Principal coordinates analysis (PCoA) revealed distinct clustering of samples from the IN4 group, indicating significant differences in their overall caecal microbial composition compared to other dietary groups (Fig. 3C). Pairwise PERMANOVA comparison showed that IN4 differed significantly from CG (R2 = 0.21, p = 0.02), CE1 (R2 = 0.24, p = 0.04), and CE4 (R2 = 0.15, p = 0.04) groups, while no significant differences were observed between other dietary groups.

**Fig. 3.**
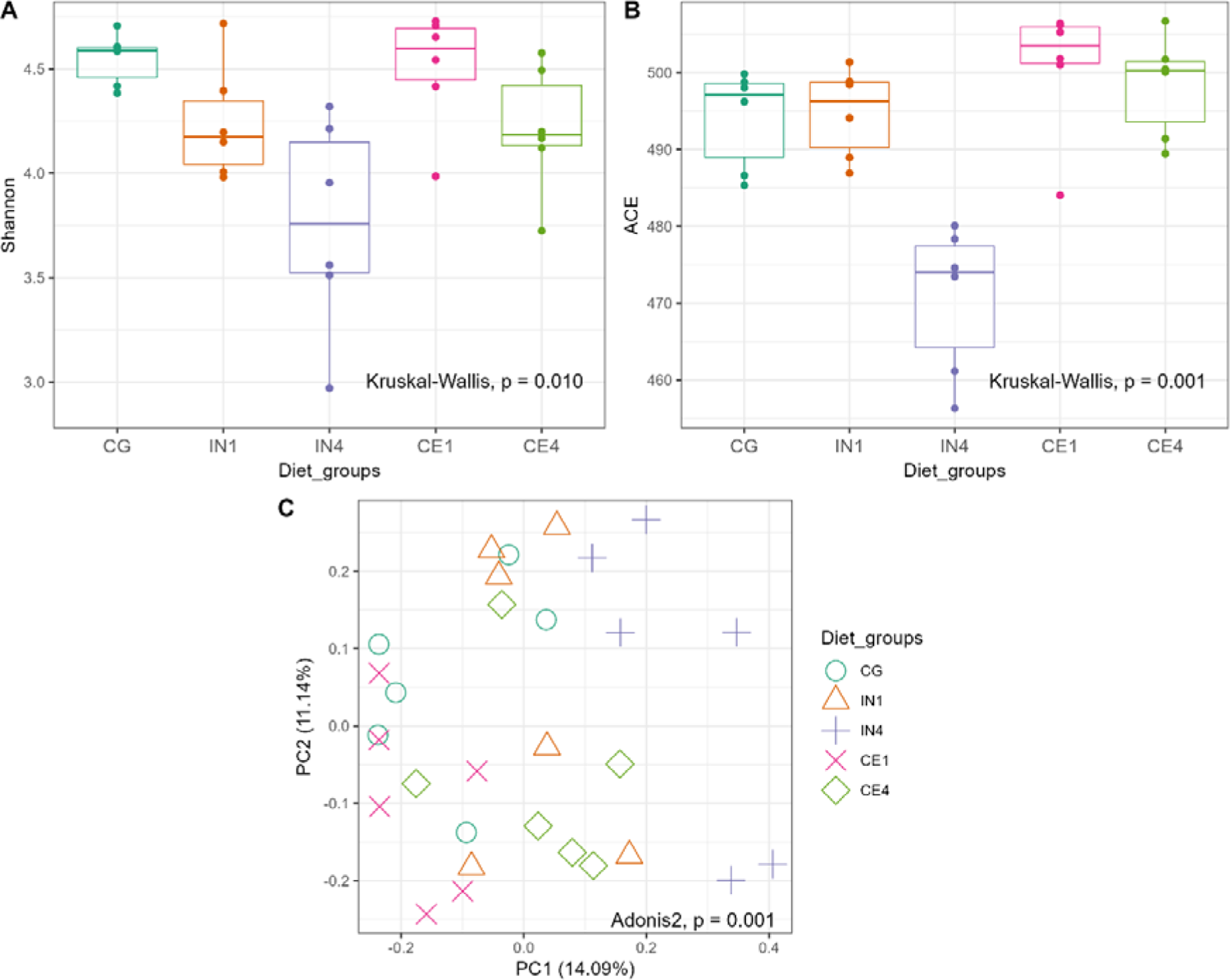
Influence of different quantities and types of dietary fibres on the diversity of caecal microbiota of chicken. Shannon (A) and ACE (B) indices showing evenness and richness of species across different dietary groups. The line inside each box represents the median value, the top and bottom lines indicate the 75th and 25th percentiles, respectively, while outliers, represented as dots, indicate values more than 1.5 times the interquartile range from the top or bottom of the box. Principal coordinate analysis (PCoA) of the caecal microbial community calculated by Bray-Curtis distance (C). Each symbol represents an individual sample, with the same symbols and colours indicating the same dietary group. Significance was defined as pL≤ 0.05. CG, Control group; IN1, 1% inulin; IN4, 4% inulin; CE1, 1% ARBOCEL; CE4, 4% ARBOCEL.

### MAGs belonging to Bacillota_A dominated the chicken caecal microbial community

We identified highly abundant MAGs in our metagenomic data (Fig. 4). All the MAGs belonged to 15 phyla, out of which Bacillota_A (64.4% ± 2.61) and Bacteroidota (13.4% ± 1.85) showed the highest relative abundance across all dietary groups (Fig. 4A). Other major phyla included Actinomycetota (5.76% ± 1.84), Bacillota_I (5.37% ± 0.48), Bacillota (4.66% ± 0.41), and Cyanobacteriota (3.39% ± 0.93). Out of 259 genera, Faecalibacterium (8.96% ± 1.46), Barnesiella (5.88% ± 1.27), and Mediterraneibacter (5.56% ± 0.58) were highly abundant across all dietary groups (Fig. 4B). At the species level, MAGs belonging to Barnesiella merdigallinarum (5.88% ± 1.27), Faecalibacterium gallistercoris (4.02% ± 0.90), Bacteroides fragilis (2.86% ± 0.55), and Bifidobacterium pullorum_B (2.77% ± 1.51) were highly abundant across all dietary groups (Fig. 4C). MAGs (n=26) that were not assigned any GTDB-TK taxonomy were grouped under Unassigned.

**Fig. 4.**
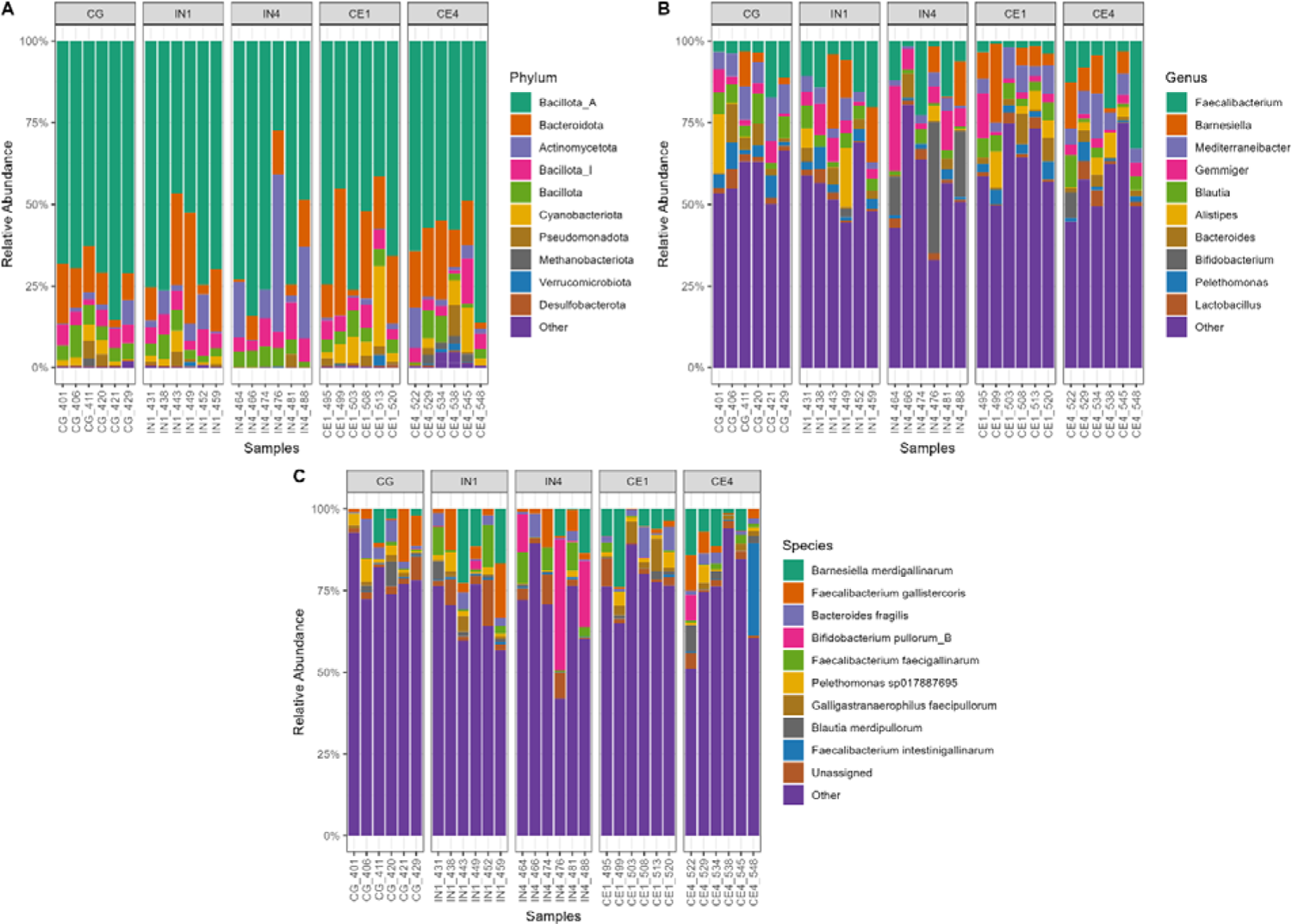
The relative abundance of caecal microbiota of chickens at phylum (A), genus (B), and species (C) levels across different dietary groups. Each column denotes a sample. Only the top 10 most abundant taxa are shown, while the remaining taxa are grouped as other. The MAGs that couldn’t be assigned any taxonomy are grouped as unassigned. CG, Control group; IN1, 1% inulin; IN4, 4% inulin; CE1, 1% ARBOCEL; CE4, 4% ARBOCEL.

### High inulin quantity exhibited a strong modulating effect on the caecal microbial community

Differential abundance analysis was performed to explore the changes in the caecal microbial community of chickens. Since high inulin supplementation significantly affected the caecal microbial diversity in our data, we focused on high dietary fibre (4%) groups for further analysis. We performed pairwise comparisons of the CG group with IN4 and CE4 groups to explore the effect of high quantities of different dietary fibres on the caecal microbial community (Fig. 5). At the phylum level, Cyanobacteriota (p < 0.001) and Bacillota_G (p < 0.001) showed significantly lower relative abundance in the IN4 group compared to CG group. At the same time, Bacillota_C (p < 0.001) and Verrucomicrobiota (p < 0.01) were significantly more abundant in the CE4 group compared to the CG group (Fig. 5A). We observed changes in the relative abundances of several genera and MAG in the IN4 group, showing a modulating effect of a high quantity of inulin on caecal microbial composition (Fig. 5B and C). At the genus level, Caproicibacterium (p < 0.001) showed significantly higher abundance in the IN4 group compared to the CG group (Fig. 5B). No significantly different genera were observed in CE4 compared to the CG group. For differential abundance analysis at the MAG level, we also included the MAGs that were not assigned any GTDB-TK taxonomy and included only those present in more than 10% of our samples. MAGs Lachnoclostridium_A pullistercoris (p < 0.001), Gemmiger sp904390925 (p < 0.001), Coprosoma intestinipullorum (p < 0.001), Catenibacillus faecavium (p < 0.001), and Caproicibacterium sp900554535 (p < 0.001) were significantly more abundant in the IN4 group compared to the CG group (Fig. 5C). In contrast, we didn’t observe any significant differences between the CG and CE4 groups at the genus and MAG species levels. The pairwise comparisons between other dietary groups are presented in Fig. S2.

**Fig. 5.**
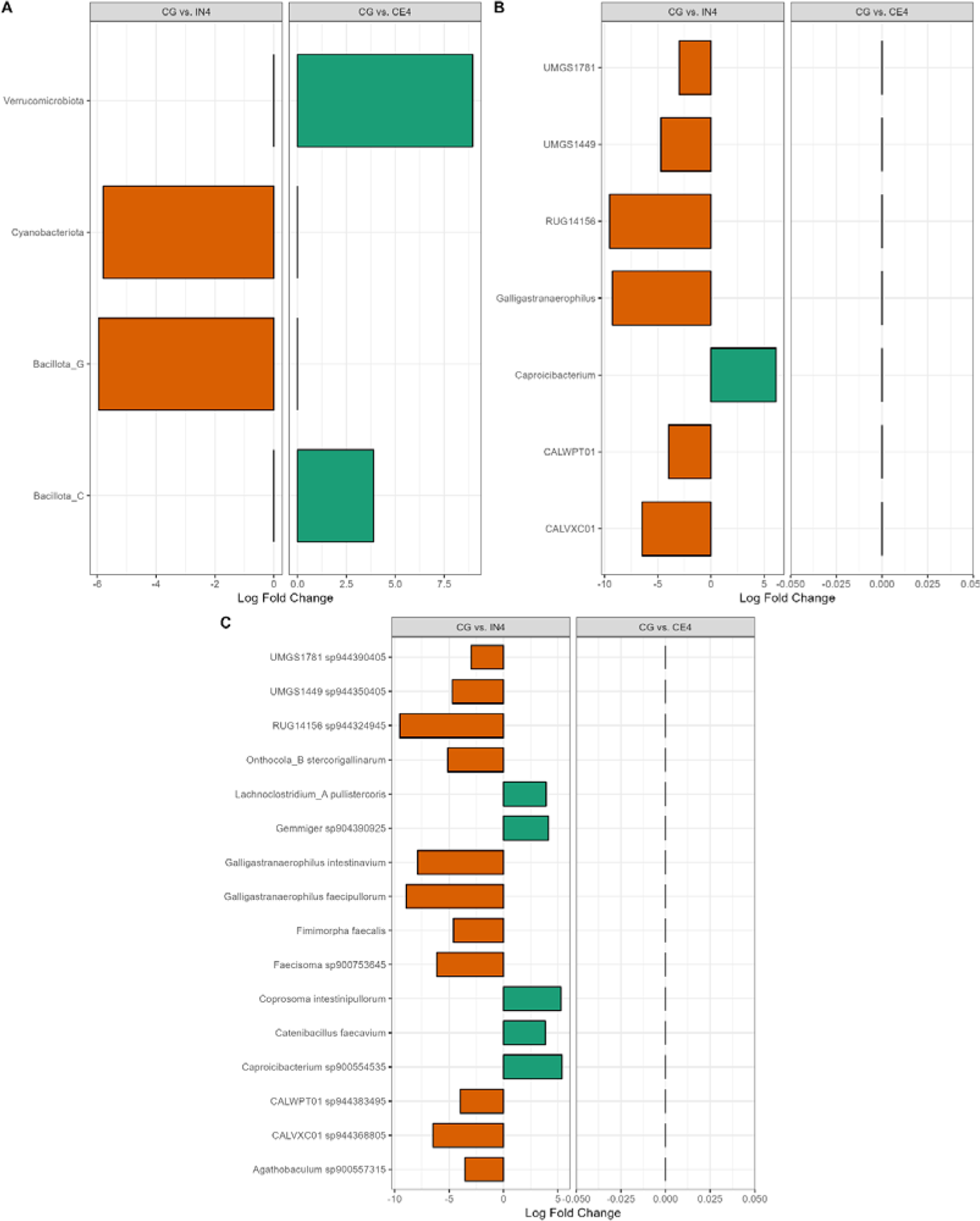
The Bar graph shows the log fold change of significantly abundant caecal microbiota at the phylum (A), genus (B), and species (C) levels in pairwise comparisons: CG vs. IN4 and CG vs. CE4. A positive log fold change value (green) indicates a higher relative abundance in the latter group, while a lower relative abundance denotes the opposite (orange). CG, Control group; IN1, 1% inulin; IN4, 4% inulin; CE1, 1% ARBOCEL; CE4, 4% ARBOCEL.

### Lower expression of carbohydrate metabolism-associated genes in the high inulin group

While metagenomics provides insights into the microbial community composition and its potential functions, metatranscriptomics provides a snapshot of actively transcribed genes, offering insight into microbial metabolic activity. We used DIAMOND-annotated metatranscriptomics data to identify alterations in microbial gene expression in response to different dietary fibres. Relative expression analysis identified DNA-directed RNA polymerase subunit beta (2.83% ± 0.11) and BMP family ABC transporter substrate-binding (1.55% ± 0.11) to be highly expressed genes across all the dietary groups (Fig. 6A, values represent mean % ± SEM). Other relatively highly expressed genes included Sn-glycerol-3-phosphate ABC transporter ATP-binding protein (1.19% ± 0.13), Stage 0 sporulation protein A (1.15% ± 0.07, D-galactose/methyl-galactoside binding periplasmic protein MglB (0.92% ± 0.15), and ATP-dependent Clp protease ATP-binding subunit (0.91% ± 0.11).

**Fig. 6.**
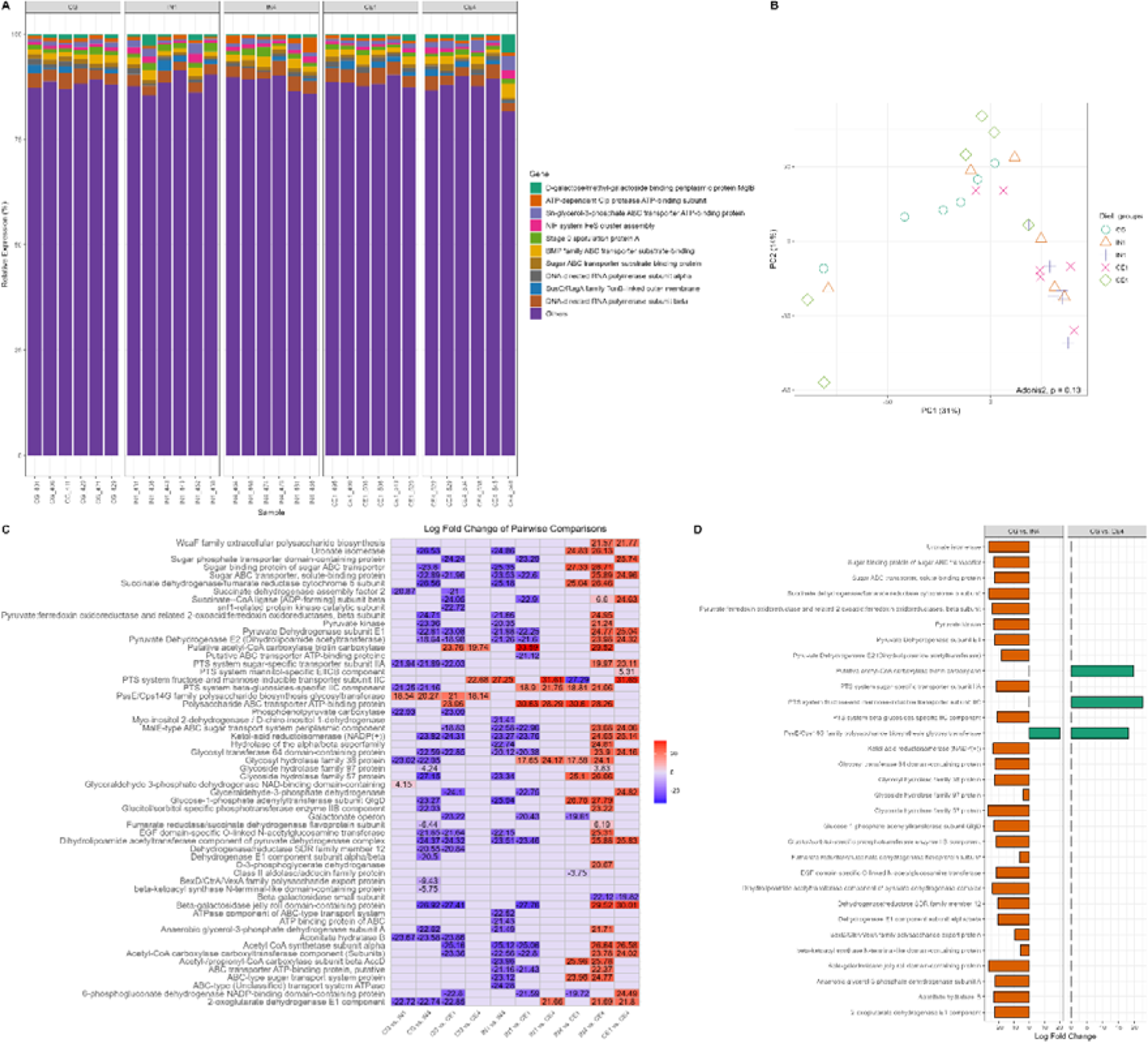
Influence of different dietary fibres on the functional profile of caecal microbiota of chickens. Principal coordinate analysis (PCoA) of the caecal microbial community calculated by Bray-Curtis distance (A). Each symbol represents an individual sample, with the same symbols and colours indicating the same dietary group. The relative gene expression of caecal microbiota of chickens across different dietary groups (B). Heatmap illustrates the log fold change of differentially expressed carbohydrate-associated genes in pairwise comparison of different dietary groups (C). The Bar graph presents a log fold change of significantly differentially expressed genes in pairwise comparisons, CG vs. IN4 and CG vs.

Differentially expressed genes (DEGs) across different dietary groups were identified to explore the effect of different types and quantities of dietary fibres on the functional profile of the caecal microbial community of chickens. A total of 546 DEGs were identified across all dietary groups using DESeq2. A heatmap based on pairwise comparison of the top 30 DEGs across different dietary groups is shown in Fig. S3. Bray–Curtis dissimilarity analysis of normalised metatranscriptomics data using the variance stabilizing transformation function revealed no significant differences in overall microbial gene expression profiles among dietary groups (Adonis2, p = 0.13). Ordination analysis further confirmed the absence of distinct clustering of samples by different dietary groups (Fig. 6B).

We specifically studied carbohydrate-associated DEGs to investigate the impact of different dietary fibres on the expression of these genes. A total of 58 carbohydrate-associated DEGs across all dietary groups were identified (Fig. 6C). We performed pairwise comparisons of DEGs from IN4 and CE4 groups with the CG group to understand the differences in microbial functional profiles in response to high quantities of different dietary fibres (Fig. 6D). Only one carbohydrate-associated DEG, i.e., PssE/Cps14G family polysaccharide biosynthesis glycosyltransferase was upregulated in the IN4 group. The upregulation of this DEG was also observed in the CE4 group, along with two other DEGs, i.e., putative acetyl-CoA carboxylase biotin carboxylase and PTS system fructose and mannose-inducible transporter subunit IIC. We didn’t find any down-regulated DEGs in the CE4 group compared to the CG group. Overall, most of the DEGs involved in glycolysis, citric acid cycle, starch and glycogen biosynthesis, and glycoside hydrolase/glycosyltransferase activity showed lower expression in the IN4 group.

CE4 (D). A positive log fold change value (green) indicates an upregulated expression in the latter group, while a downregulated expression denotes the opposite (orange). Significance was declared at p ≤10.05. CG, Control group; IN1, 1% inulin; IN4, 4% inulin; CE1, 1% ARBOCEL; CE4, 4% ARBOCEL.

### High inulin quantity upregulated CAZyme families GH1 and GH32 identified using dbCAN3

We next investigated carbohydrate-active enzymes (CAZYmes) genes, as they play a crucial role in enabling the gut microbiota to degrade and utilize dietary fibres. We identified a total of 119,269 CAZyme coding genes in our metatranscriptomics data using the dbCAN3 database. Glycoside hydrolases (GH) and glycosyltransferases (GT) were the most highly expressed CAZyme classes, accounting for 51.2% and 33.5%, respectively (Table S1). GH is known to cleave glycosidic bonds to break down carbohydrates, while GT is involved in the formation of new glycosidic bonds to form or modify carbohydrate structures. Carbohydrate esterases (CE) class represented 7.7% and carbohydrate binding modules (CBM) represented 6.3% in our data. Some of the reads (0.17%) showed similarity to CAZymes, but could not be assigned to any known CAZyme class. This might be due to their low similarity to characterized CAZymes or limitations in the current CAZyme databases. Out of 552 CAZyme families identified in our data, GT2 (11.6% ± 0.21), GH13 (7.38% ± 0.12), GT4 (5.99% ± 0.14), and GH3 (4.89% ± 0.12) were highly expressed across all dietary groups (Fig. 7A, values represent mean % ± SEM).

**Fig. 7.**
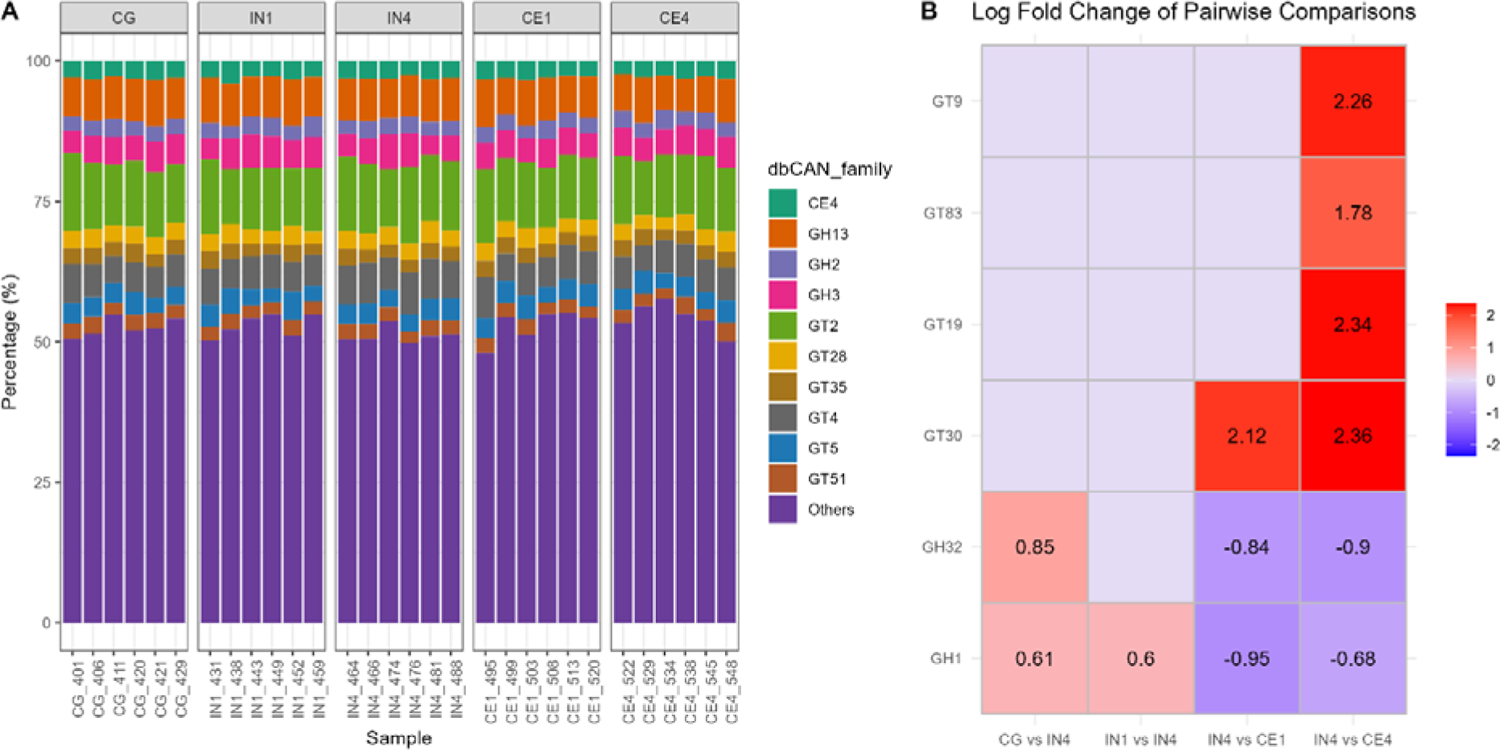
Influence of different dietary fibres on the CAZyme encoding genes across all the dietary groups. The bar graph shows highly expressed CAZyme families in the chicken caecal microbial community. Heatmap presents the log fold change of differentially expressed CAZyme families in pairwise comparison of different dietary groups, highlighting only significant results. Significance was declared at p ≤10.05. CG, Control group; IN1, 1% inulin; IN4, 4% inulin; CE1, 1% ARBOCEL; CE4, 4% ARBOCEL.

We then used the GH and GT classes to perform differential expression analysis due to their high relative abundance in our data and key roles in carbohydrate metabolism. Adonis2 analysis based on Bray-Curtis dissimilarity showed significant differences in GH (Adonis2, p = 0.001) and the GT (Adonis2, p = 0.01) classes. Ordination analysis showed separate clustering of CG and IN4 groups in GH class, while no clustering was observed in the case of GT class (Fig. S4). Pairwise PERMANOVA comparison for the GH class showed significant differences in the IN4 group compared to CG (R2 = 0.21, p = 0.03) and CE1 (R2 = 0.24, p = 0.01) groups, while no significant differences were observed between other dietary groups. We didn’t find any pairwise significant difference between different dietary groups in the GT class.

We then used DESeq2 to perform pairwise comparisons between different dietary groups to identify significantly differentially expressed CAZyme families. A few of our pairwise dietary group comparisons showed significant differences. The upregulated expression of GH32 and GH1 families was found in the IN4 group compared to the CG group. The GH32 family enzymes, including inulinases and levanases, specifically break down inulin-type fructans into fermentable sugars. In contrast, the GH1 family includes β-glucosidases and β-galactosidases, which broadly hydrolyse β-linked oligosaccharides and glycosides in plant cell walls. The GT families, including GT9, GT83, GT19, and GT30, showed upregulation, and GH families, including GH32 and GH1, showed downregulation in the CE4 group compared to the IN4 group (Fig. 7B). No significant differences were observed between CE4 and CG groups for either family. These GT families are involved in the biosynthesis of various structural and functional glycoconjugates like lipopolysaccharides, exopolysaccharides, and glycoproteins. They transfer sugar moieties to various acceptors, supporting cell wall integrity, signalling, and interactions within microbial communities. Overall, the results suggest that the caecal microbiota in the CE4 group were more engaged in anabolic processes, whereas those in the IN4 group were more involved in sugar hydrolysis and degradation processes.

### MAGs showed a similar expression profile but different expression levels in different dietary groups

We used MAGs and already identified significantly expressed CAZymes and carbohydrate-associated DEGs from our metatranscriptomics data to link them to their corresponding MAGs to find to what degree MAG genes are being expressed in different samples, thereby elucidating their roles in dietary fibre digestion. We compared the IN4 and CE4 groups with the CG group by using the significantly different DEGs and CAZymes data from our metatranscriptomics analysis (Fig. 8 A-B).

**Figure 8:**
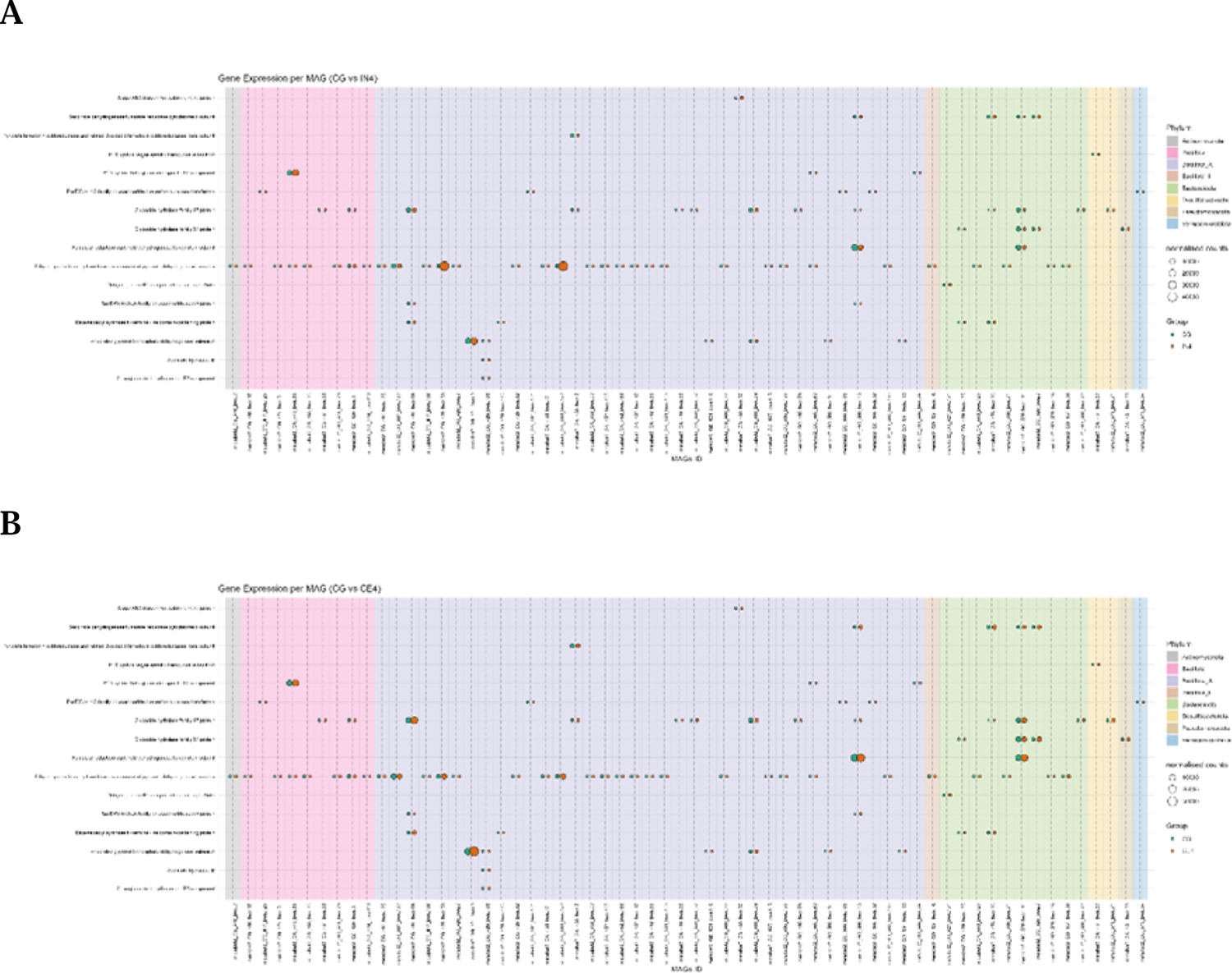
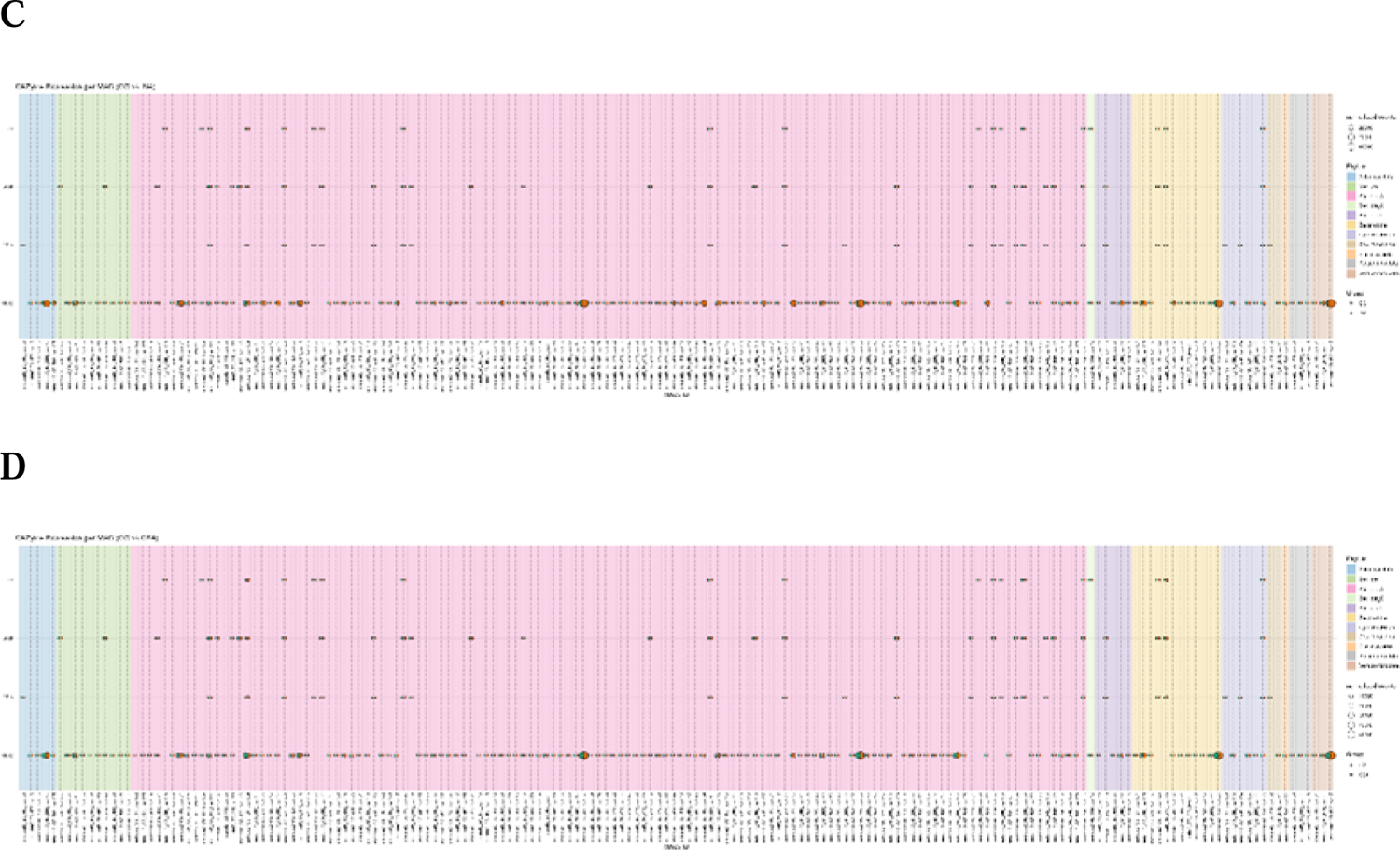
Bubble plots show the different expression levels of significantly differentially expressed carbohydrate-associated genes (A and B) and CAZymes (C and D) by different MAGs in IN4 and CE4 compared to the CG group. Bubble size represents normalised counts per gene per MAG across different dietary groups. Background colour indicates the phylum to which each MAG belongs. CG, Control group; IN1, 1% inulin; IN4, 4% inulin; CE1, 1% ARBOCEL; CE4, 4% ARBOCEL.

Out of 514 MAGs, 340 MAGs showed the expression of DEGs. We then filtered out genes which were expressed by 55% of the MAGs, including, Glucose-1-phosphate adenylyltransferase subunit GlgD, Pyruvate kinase, and Uronate isomerase and ended up with 62 MAGs. Most of the MAGs expressing DEGs belonged to Bacillota_A. Overall, the distribution of normalised gene counts across MAGs looked similar in CG vs. IN4 and CG vs. CE4 comparisons. However, group-specific differences in read counts were observed across dietary treatments for some MAGs. Gene counts for succinate dehydrogenase/fumarate reductase cytochrome b subunit by MAG metabat2_GC_520_bwa.111 (p_Bacteroidota;s_Alistipes megaguti) were lower in IN4 compared to the CG group, while no apparent difference was noted between the CE4 and CG groups. Higher gene counts for PssE/Cps14G family polysaccharide biosynthesis glycosyltransferase were observed in the CG group compared to IN4 and CE4 groups by different MAGs, including metabat2_GA_431_bwa.101 (p_Bacillota_A;s_Flemingibacterium merdigallinarum), metabat2_GA_452_bwa.105 (p_Bacillota;s_Scatomorpha sp900759385), and metabat2_GC_508_bwa.32 (p_Bacillota_A;s_Scatomorpha sp900545405), while an opposite trend was observed by MAG metabat2_CG_411_bwa.125 (p_Bacillota;s_Limosilactobacillus oris). These results indicate specific functions performed by different MAGs in the digestion of dietary fibre in the caeca of chicken. The details of other DEGs showing higher gene counts in IN4 and CE4 groups compared to the CG group are presented in Table 1.

We then analysed the gene counts for CAZymes by the MAGs (Fig. 8C-D). For this, we used CAZymes identified as significantly differentially expressed in our metatranscriptomics analysis to understand the contribution of individual MAGs in their expression. Overall, a total of 349 MAGs were involved in the expression of significantly differentially expressed CAZymes across dietary groups. GH1 and GT83 were expressed by ≥80% of all the MAGs, which were removed for downstream analysis, leaving 176 MAGs for further study. The distribution of gene counts for CAZymes, including GH32, GT19, GT30, and GT9, was visualised in CG vs. IN4 and CG vs. CE4 comparisons. Just like the DEGs, the overall distribution of normalised CAZyme gene counts was similar across all dietary groups, but group-specific differences were observed for some MAGs. Higher gene counts of the GH32 family were observed in IN4 and CE4 groups compared to the CG group by different MAGs. The details and taxonomies of MAGs are presented in Table S5. The gene counts for GH32 were lower in IN4 and higher in CE4 group compared to CG group by metabat2_CG_420_bwa.139 (p_Bacillota_A;s_Scatosoma pullicola). We observed higher gene counts for GT30 in the CE4 group compared to the CG group by MAGs (Table S5), however, no differences were observed between the IN4 and CG groups.

### Dietary fibres showed minimal influence on the carbohydrate-associated protein expression of caecal microbiota

We performed metaproteomics analysis to understand which proteins were produced by caecal microbiota in response to different dietary fibres. A total of 48554 unique proteins were initially identified across all dietary groups, which was reduced to 6996 after applying a filtering step to retain only proteins quantified in all replicates of at least one group. We identified 63 differentially expressed proteins (DEPs) through pairwise comparison between different dietary groups, out of which only five DEPs were carbohydrate-associated (Fig. S5). Pairwise comparison of IN4 and CE4 with the CG group showed higher relative expression of glyceraldehyde-3-phosphate dehydrogenase A and lower relative expression of alpha-galactosidase in the CE4 group, while higher relative expression of pyruvate, phosphate dikinase, and lower relative expression of pyruvate oxidase were observed in the IN4 group (Fig. 9).

**Fig. 9.**
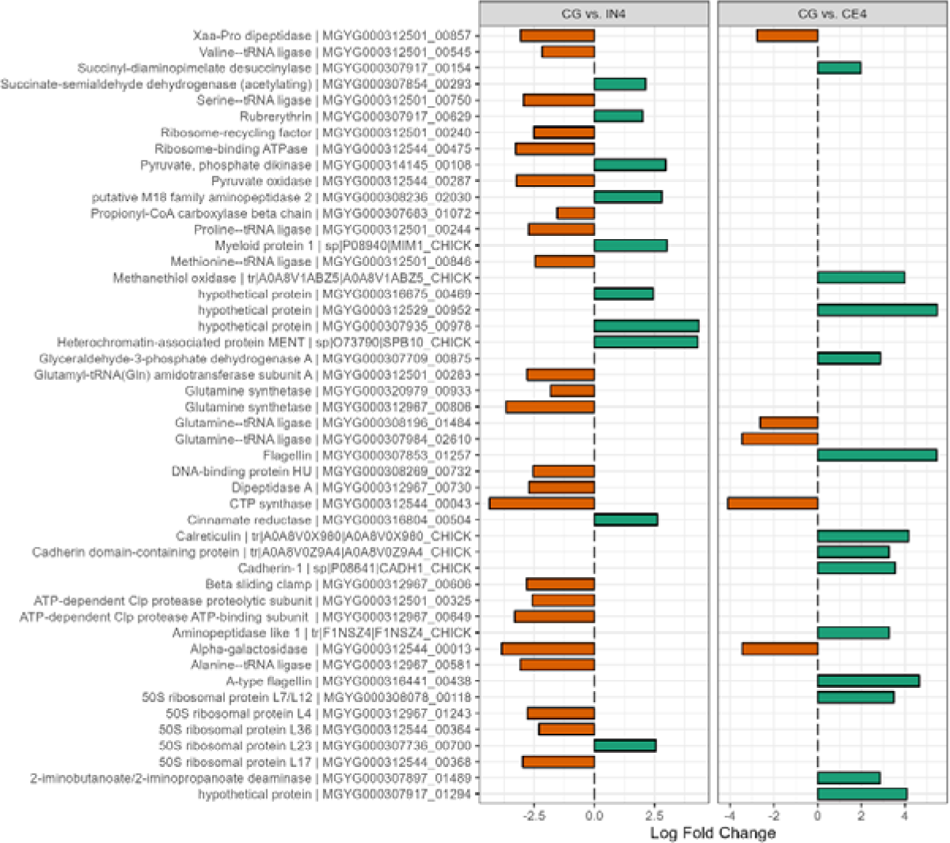
The Bar graph shows the log fold change of significantly differentially expressed proteins in pairwise comparisons, CG vs. IN4 and CG vs. CE4. A positive log fold change value (green) indicates an upregulated expression in the latter group, while a downregulated expression denotes the opposite (orange). Significance was declared at p ≤10.05. CG, Control group; IN1, 1% inulin; IN4, 4% inulin; CE1, 1% ARBOCEL; CE4, 4% ARBOCEL.

### Dietary fibres showed minimal effect on the expression of immune-associated genes in the host caecal epithelium

Dietary fibre is known to influence the immune system of the host through gut microbiota. Therefore, we explored immune-associated DEGs across all dietary groups (Table S2). We did not identify any DEGs related to the immune system in the IN1 and CE1 groups compared to the CG group. However, we identified downregulations of two immune-associated DEGs i.e., CCR8L and S100A10, in the IN4 groups, while three immune-associated DEGs, i.e., GPC3, FOXN1, and EFNA2, showed upregulations in the CE4 group compared to the CG group. The majority of DEGs lacked functional annotation or were assigned to hypothetical proteins, making it challenging to interpret the results.

## Discussion

Dietary fibre is incorporated into poultry rations as an alternative feed resource to improve gut health, reduce feed-food competition, and promote more sustainable poultry production. Dietary fibre was once thought to be anti-nutritional (Pérez-Jiménez 2024); however, its beneficial effects on supporting the immune system, modulating gut microbiota, and improving digestive efficiency have now been acknowledged (Makki et al. 2018; Jha & Mishra 2021). In the chicken’s digestive system, the caecum is the main site of microbial fermentation, harbouring a complex and dynamic microbial ecosystem. The caecal microbial community plays an important role in the digestion of dietary fibre to extract energy from otherwise indigestible components. Among the various fibre types, Inulin and cellulose represent two contrasting fibre sources with distinct physicochemical and fermentative characteristics. Inulin is a type of fructan that is a well-documented prebiotic and promotes the growth of beneficial gut microbiota (Anand et al. 2025). Cellulose is a complex, insoluble polysaccharide that enhances gut health and improves nutrient utilization by supporting microbial fermentation (Yasmeen & Ahmad 2025). Understanding how these dietary fibre influences the composition and function of gut microbiota is key to improving poultry production through targeted microbiota manipulation. In this study, we examined the effects of inulin as soluble and cellulose as insoluble dietary fibres on the composition and functional potential of caecal microbiota in broiler chickens using a multi-omics approach.

We compared the taxonomic assignments of metagenomic and metatranscriptomic reads in response to different dietary fibres to identify which taxa are abundant and which are functionally active. Among the top ten genera identified, we observed higher read counts of Bifidobacterium in metagenomic data, which Agathobaculum replaced in metatranscriptomics data. This difference highlights that high microbial abundance does not necessarily reflect metabolic activity and functionally active taxa might be underrepresented in DNA-based profiles. Another study reported similar findings, where several low-abundance genera showed metabolic activity in response to dietary fibre supplementation (Xia et al. 2021). These findings highlight the importance of integrating metatranscriptomic analyses to complement taxonomic profiling and gain a more comprehensive understanding of the functional dynamics of the gut microbial community.

In this study, high inulin quantity significantly reduced the microbial diversity, which is inconsistent with previous reports showing that inulin increased microbial diversity (Chen et al. 2025). This discrepancy might be attributed to the quantity of inulin used, as our study used 4% inulin concentration compared to 1.2% used in the previous study. Based on these distinct changes, we focused on the high fibre groups to better understand how increased dietary fibre shaped the caecal microbial community in chicken. We found that high inulin had a stronger influence on caecal microbial community than high cellulose, with significant variations observed at phylum, genus, and MAG levels. Previous studies reported an increase in the genus Bifidobacterium and Lactobacillus in response to inulin (Song et al. 2020; Abdelqader et al. 2013; Liu et al. 2018). However, in our study, Caproicibacterium showed significantly higher abundance in the IN4 group. Although Bifidobacterium and Lactobacillus were highly abundant in the inulin groups, they didn’t show significant differences compared to the control group, consistent with previous findings (Xia et al. 2021). This suggests that high inulin favoured the growth of alternative fibre-degrading or fatty acid-producing taxa such as Caproicibacterium (Zeng et al. 2024), which might be able to utilize inulin-derived substrates under high fibre conditions. On the other hand, the influence of high cellulose quantity was minimal on microbial composition, with significant variations primarily limited to the phylum level only. This aligns with observations reported in a previous study where lignocellulose supplementation didn’t modify the ileal microbial composition in young broiler chickens (Farkas et al. 2025). These differences may be due to the recalcitrant nature of cellulose (Feng et al. 2025), which makes it challenging for microbes to break it down, compared to the more fermentable and accessible inulin. These findings highlight the significance of fibre type and its structure, suggesting that soluble fibre might be more effective in modulating the caecal microbial community of broilers.

While studying the microbial functional profile, we found a decreased expression of genes associated with core metabolic processes, such as glycolysis, citric acid cycle, starch, and glycogen biosynthesis in the IN4 group. Similar findings were reported in the caecal microbiota of Salmonella-infected chickens supplemented with inulin (Song et al. 2020). Moreover, lower expression of pyruvate oxidase and higher expression of pyruvate, phosphate dikinase (PPDK) proteins were also noted in this study. Both of these enzymes are involved in pyruvate metabolism and work through distinct metabolic pathways.

Pyruvate oxidase is involved in pyruvate decarboxylation and steers the metabolic pathway toward acetate production (Abdel-Hamid et al. 2001), while PPDK is associated with gluconeogenesis and energy conservation by reversing glycolytic pathways under anaerobic conditions (Chastain et al. 2011). These shifts suggest a broader metabolic adaptation by the microbial community, moving away from central energy metabolism to more specialized fermentative processes tailored to the inulin-rich environment. These functional changes were further supported by significantly higher expression of GH32 and GH1 families in the IN4 group. GH32 encodes inulinase, an enzyme specifically involved in the breakdown of inulin-type fructans into fermentable sugars (Pouyez et al. 2012). GH1 encodes β-glucosidases, which cleave glycosidic bonds between glucose units in oligosaccharides and play a crucial role in the final steps of carbohydrate degradation (Kaenying et al. 2023). Moreover, the majority of our MAGs showed expression of the GH1 gene, indicating its functional importance within the caecal microbial community. The increased expression of these CAZyme genes might suggest that while microbes prefer readily available inulin by expressing GH32, they are also showing broader adaptation to secondary β-glucosidic substrates released during microbial fermentation.

On the other hand, the functional changes in caecal microbiota in response to high cellulose quantity were more subtle. We observed higher expression of three carbohydrate-associated genes linked to sugar transport, fatty acid synthesis, and polysaccharide biosynthesis. Additionally, higher expression of the enzyme glyceraldehyde-3-phosphate dehydrogenase A was observed in response to high cellulose quantity. It is a key glycolytic enzyme (Nicholls et al. 2012), indicating increased glycolytic activity to support basal energy demand with limited fermentable substrates. These findings suggest that microbes in the CE4 group preferred corn/soy components, as cellulose is challenging to break down and extract energy through glycolysis and the TCA cycle for fatty acid production, and perform biosynthetic activity to maintain cellular structure. These findings highlight the distinct functional strategies employed by caecal microbes in response to different dietary fibre types. While inulin promotes active sugar degradation and enzyme expression, cellulose tends to elicit more structural or regulatory responses, likely due to its recalcitrance and lower fermentability. These underscore the importance of fibre type and quantity in shaping not only microbial composition but also metabolic function within the gut ecosystem.

This study is the first to use multi-omics approaches to capture both microbial composition and their functions at the transcript and protein levels, in relation to fibre fermentation in chickens. The findings provide insights into how specific fibre types influence not only microbial composition but also their active metabolic roles within the gut. Ultimately, this research lays the groundwork for developing targeted, fibre-based dietary strategies to enhance gut health, nutrient utilisation, and productivity in poultry production systems.

While our integrated multi-omics approach offers valuable insights into the composition and functional activity of the caecal microbiota, several limitations should be acknowledged. First, the study was based on caecal samples collected at a single time point, which may not reflect temporal changes in microbial dynamics. Second, both shotgun metagenomic and metatranscriptomic analyses rely on existing reference databases, which may underrepresent certain microbial taxa or functional genes, potentially limiting resolution. Finally, although we characterised microbial function at the transcript and protein levels, we did not directly assess downstream functional outcomes such as short-chain fatty acid production or host physiological responses. These aspects warrant further investigation to fully elucidate the implications of dietary fibre interventions in poultry.

## Conclusions

This study offers comprehensive insights into how different types of dietary fibre — soluble inulin and insoluble cellulose — modulate the composition and functional activity of the caecal microbiota in broiler chickens. Using a multi-omics approach, we found that high inulin quantity significantly altered microbial diversity, enhanced the expression of GH32 genes, and prompted metabolic shifts away from central energy pathways, indicating microbial adaptation to readily fermentable fibre. In contrast, high cellulose had a minimal influence on microbial community and enhanced glycolytic activity to support the basic energy needs of microbes with limited fermentable substrate. These findings highlight the importance of fibre type and quantity in shaping gut microbial community and their functions, with potential implications for designing strategies aimed at modulating microbiota to improve feed efficiency, nutrient utilization, and overall poultry health.

## Methods

### Experimental Design and Sampling

The study was approved by the Roslin Institute Animal Welfare and Ethical Review Board. Chickens were housed in premises licensed under a UK Home Office Establishment License within the terms of the UK Home Office Animals (Scientific Procedures) Act 1986. Housing and husbandry complied with the Code of Practice for Housing and Care of Animals Bred, Supplied or Used for Scientific Purposes.

### Experimental Design

The study was conducted at the National Avian Research Facility, The Roslin Institute, University of Edinburgh, UK. Ninety chickens (1-day-old male Ross 308s broilers) sourced from a commercial hatchery (PD Hook LTD, Cote, Bampton) were divided into five dietary groups, each consisting of six pens with three birds per pen in a completely randomized design. The five dietary groups were (1) control diet (CG): received a standard corn/soya-based diet with no added fibre; (2) grower A (IN1): received corn/soya-based diet supplemented with 1% inulin during both the starter and the grower phase; (3) grower B (IN4): received corn/soya-based diet supplemented with 1% inulin during starter phase and 4% inulin during the grower phase; (4) grower C (CE1): received corn/soya-based diet supplemented with 1% ARBOCEL during both the starter and grower phase; (5) grower D (CE4): received corn/soya-based diet supplemented with 1% ARBOCEL during starter phase and 4% ARBOCEL during grower phase. All birds had free access to feed and water throughout the study period. The details of raw ingredients and nutritional composition of starter and grower diets across different dietary groups are presented in Table S3. At day 35, one bird from each pen was randomly selected and humanely killed via cervical dislocation for post-mortem caecal sampling for multi-omic analysis. The confirmation of death was performed through cessation of circulation.

The caecal contents (n=30; 6 birds/group) from both caeca were removed and mixed. The homogenised content was then divided into 3 tubes: one tube containing 750µl RNAlater for RNA analysis, and the other two separate tubes designated for DNA and protein analysis. Moreover, a section of mid-caecal epithelium was also excised and stored in a tube containing 750µl RNAlater. The RNAlater-containing tubes were placed at 4°C for 24 hours to allow RNAlater to permeate tissue and stabilise RNA by inhibiting RNase activity. The tubes were later stored at −80°C until RNA isolation. The tubes for DNA and protein analysis were immediately snap-frozen on dry ice and stored at −80°C until further analysis.

### Nucleic acid extraction and sequencing

Genomic DNA (n=30) and total RNA (n=30) were extracted from each homogenised caecal content using QIAamp PowerFecal Pro DNA Kit (QIAGEN®) and RNeasy PowerMicrobiome Kit (QIAGEN®) according to the manufacturer’s instructions, respectively. Extracted DNA samples were treated with RNase Cocktail™ Enzyme Mix (Thermo Fisher) at a 1:20 volume ratio and incubated at room temperature for 15-30 minutes to remove RNA contamination. Additionally, total RNA was isolated from caecal mid-epithelium using RNeasy® Mini kit (QIAGEN®) with a minor modification. During the initial pulverisation step, one 5mm stainless steel bead was added to each sample to enhance tissue lysis. DNA and RNA samples were purified using AMPure XP Beads (Beckman Coulter) at a modified 1:1 volume ratio, followed by the remaining steps as per the manufacturer’s instructions. For RNA samples, the beads were pre-treated with RNaseOUT™ Recombinant Ribonuclease Inhibitor (Thermo Fisher) according to the manufacturer’s instructions before the purification step. The quantity and quality of extracted nucleic acid were checked using Qubit® 4.0 (Invitrogen Life Technologies, Carlsbad, CA, USA) and agarose gel electrophoresis.

The shotgun metagenomics (DNA from caecal content), metatranscriptomics (total RNA from caecal content), and eukaryotic mRNA (total RNA from caecal mid-epithelium) sequencing of purified samples were performed by Novogene Corporation Inc using a NovaSeq Illumina platform, producing paired-end 150 bp reads. Metaproteomics analysis on caecal content was conducted by the Proteomics & Metabolomics Facility at The Roslin Institute, University of Edinburgh.

### Metagenomics data analysis

The shotgun metagenomics sequencing generated 12Gb of raw reads per sample. The adapters and low-quality reads were removed using fastp (v0.23.4), and reads aligned to the chicken genome (release-111, bGalGal1.mat.broiler.GRCg7b, database downloaded 6 June 2024) were removed using BWA MEM (v0.7.18) (Chen 2023; Li 2013). A de novo assembler, MEGAHIT (v.1.2.9), was used to generate single assemblies for each sample using the options --continue --kmin-1pass --k-list 27,37,47,57,67,77,87 --min-contig-len 1000 parameters (Li et al. 2015). The assembled contigs were indexed, and reads were mapped to assemblies using BWA MEM (v0.7.18). The resulting alignments were sorted and converted to BAM format using SAMtools-sort (v1.20), and summary statistics were generated using the command SAMtools-flagstat (Li et al. 2009). Coverage depth per contig was calculated using jgi_summarize_bam_contig_depths command from MetaBAT2 (v2.15) (Kang et al. 2019). Metagenomic binning was subsequently executed using MetaBAT2, and the quality of the resulting Metagenome Assembled Genomes (MAGs) was assessed with CheckM2 (v1.0.1) (Chklovski et al. 2024). The MAGs were dereplicated using dRep (v3.5.0) at 95% and 99% of Average Nucleotide Identity (ANI), and MAGs having completeness ≥80% and contamination ≤10% were filtered. However, MAGs dereplicated at 95% were used for downstream analysis to capture species-level diversity and reduce redundancy (Evans & Denef 2020). The taxonomic assignment of the MAGs was carried out with GTDB-tk using the classify_wf function with default parameters (v2.4.0, database downloaded 5 August 2024), and abundance estimation was performed using CoverM with --min-read-percent-identity 95 and --min-read-aligned-percent 85 parameters (v0.7.0) (Chaumeil et al. 2022; Aroney et al. 2025). The phylogenetic tree was constructed using phylophlan (v3.1.1) and rerooted using FigTree (v1.4.3) at the branch between the archaeal and bacterial MAGs. The rerooted tree was then visualized using iTOL v6 (https://itol.embl.de/). The phyloseq (v1.42.0) package in RStudio (v4.2.3) was used to calculate alpha (Shannon and ACE indices) and beta (Bray-Curtis distance) diversities based on TPM (Transcripts Per Million) values generated by CoverM (McMurdie & Holmes 2013). The TPM values were transformed into raw count equivalents to make them suitable for ANCOM-BC2 (v2.0.3) (Lin & Peddada 2020). Differential pairwise comparisons were then performed at the phylum, genus, and species levels using the Dunn test in ANCOM-BC2.

### Metatranscriptomics data analysis

After assessing the quality of raw reads using fastQC (v0.11.7), FastP (v0.23.4) was used to remove poor-quality sequences and adaptors, while SortMeRNA (v4.3.6) was used to eliminate rRNA reads (Kopylova et al. 2012). The clean reads were mapped to the chicken transcriptome (release-111, bGalGal1.mat.broiler.GRCg7b, database downloaded 31 May 2024) from Ensembl (https://www.ensembl.org/index.html) using BWA MEM (v0.7.18) to remove host contamination, and SAMtools was used to obtain the unmapped reads (Danecek et al. 2021). Then, Megahit (v1.2.9) was utilised to generate single-sample assemblies. Unlike the metagenomic analyses, the minimum length was adjusted to 300bp to capture smaller mRNAs. After indexing assemblies and mapping reads to assemblies using BWA MEM and generating summary statistics using SAMtools-flagstat, the feature counts within the coding DNA sequence region were quantified with HTSeq-count (v2.0.5) (Anders et al. 2015).

MetaGeneMark (v.3.38) was used to identify coding regions from the assemblies, and Seqkit (v2.8.2) was used to produce statistics for those coding regions (Gemayel et al. 2022; Shen et al. 2016). The DIAMOND (v2.1.8) database, built from UniRef90 FASTA (database download 8^th^ July 2024) and NCBI taxonomy files (download 9 July 2024), was used to assign functional information to the coding regions (Suzek et al. 2015). The annotation results from DIAMOND and the feature count results from the HTSeq were used for downstream analysis. DIAMOND annotated genes were filtered based on bitscore >50, and were merged with HTSeq counts based on gene ID after removing any redundancy. Counts for genes encoding the same gene product were aggregated to reflect functional shifts in microbial communities, and the resulting dataset was used for expression analysis. After pre-filtering the rows with counts ≥ 10 in a minimum of 3 samples, the remaining counts were normalised using the variance stabilizing transformation (VST) function implemented in DESeq2 (v1.38.3). The relative and differential expression analysis across different diet groups was conducted using DESeq2, and gene products with p≤0.001 were included in the further analysis.

CAZyme annotation was performed by dbCAN3 (database built on 11 July 2024) on coding sequences predicted by MetaGeneMark (Zheng et al. 2023). The dbCAN3 results included annotation from HMMER, dbCAN_sub, and DIAMOND databases. Enzymes annotated by at least two databases were included to explore the relative expression of CAZyme types and families. Moreover, GH and GT families were selected for differential expression analysis using DESeq2 (v1.38.3) due to their higher expression in this study and key role in carbohydrate metabolism.

### Linking MAGs to metatranscriptomics gene expression

The protein-coding regions within the MAGs were predicted using Prodigal (v2.6.3) and annotated using DIAMOND BLASTp (Hyatt et al. 2010). The FASTA files of each MAG were indexed using BWA index, and quality-filtered metatranscriptomics reads from each sample were aligned to each MAG using BWA MEM. The mapped reads were used to generate BAM files using SAMtools-sort. Depth files were created to understand how many reads were mapped to each gene in MAGs using SAMtools-flagstat. Finally, HTSeq-count was used to generate gene counts by counting RNA reads aligned to annotated genes in MAGs using BAM files and the corresponding .gff annotation files generated by Prodigal.

After merging gene counts with functional annotations from DIAMOND, read counts were normalised using estimateSizeFactors function of DESeq2. Genes and CAZymes that were significantly differentially expressed in our study were retained to compare their expression in IN4 and CE4 groups with the CG group in each MAG.

### Kraken2 analysis

A custom Kraken database was built (25 June 2024) using microbial RefSeq genomes from NCBI (including bacteria, archaea, plasmids, viruses, fungi, plants, and protozoa), the chicken genome (bGalGal1.mat.broiler.GRCg7b), and GenBank assemblies from the NCBI BioProjects PRJNA715658, PRJEB64517, PRJEB33338, PRJNA543206, PRJNA377666, PRJNA668258, PRJEB57055 (data downloaded 16 June 2024). Quality-controlled reads from metagenomic and metatranscriptomic sequencing were classified at the kingdom, phylum, and genus levels using Kraken2 (v2.1.3) (Lu et al. 2017; Wood et al. 2019).

### Metaproteomics data analysis

Faecal contents were homogenised in an extraction buffer comprising 5% SDS, 6 M urea, and 50 mM triethylammonium bicarbonate (TEAB), pH 8.5, at a sample-to-buffer ratio of 1:10 (w/v). Homogenisation was performed using a Precellys homogeniser at 5000 × g for 20 seconds in a ceramic bead vial (Precellys Lysing Kit, Tissue Homogenising CK Mix). The homogenates were then centrifuged at 16,000 × g for 10 minutes, and the supernatant was transferred to low-binding protein vials. Samples were subsequently sonicated using a Bioruptor Pico Sonicator (Diagenode) for 10 cycles (30 seconds on / 30 seconds off per cycle). Following sonication, samples were centrifuged at 16,000 × g for 10 minutes. The resulting supernatant was collected, and protein concentration was determined using a BCA assay.

Tryptic digestion was carried out using S-Trap microcolumns (Protifi, USA), following the manufacturer’s protocol with minor modifications. In brief, 20 µg of protein in extraction buffer was reduced with 10 mM dithiothreitol at 37°C for 1 hour and alkylated with 18.75 mM iodoacetamide at room temperature for 35 minutes in dark conditions. Alkylation was quenched by adding phosphoric acid to a final concentration of 1.25%, followed by the addition of six volumes of binding buffer (90% methanol in 100 mM TEAB). After gentle vortexing, the protein suspension was loaded onto an S-Trap microcolumn and centrifuged at 4000 × g for 1 minute. The column was then washed three times with 150 µL of binding buffer, each with a spin at 4000 x g for 1 minute. Proteolytic digestion was performed by adding 20 µL of digestion buffer (1 µg trypsin in 50 mM TEAB) and incubating the column at 47°C for 2 hours. Peptides were sequentially eluted using 40 µL each of: (1) 50 mM TEAB, (2) 0.1% formic acid in water, and (3) 0.1% formic acid in 50% acetonitrile. Eluted peptides were pooled, cleaned using C18 stage tips, and dried under a vacuum desiccator.

Purified peptides were separated over a 70-minute gradient on an Aurora 25 cm column (IonOpticks, Australia) using an UltiMate RSLCnano LC system (Thermo Fisher Scientific) coupled to a timsTOF HT mass spectrometer via a CaptiveSpray ionisation source. The LC gradient was delivered at a flow rate of 200 nL/min, with a post-run washout step at 500 nL/min. Column temperature was maintained at 50°C. Data-dependent acquisition (DDA-PASEF) was employed, with full MS scans acquired from 100-1700 m/z and ion mobility ranging from 1.45 to 0.65 Vs/cm² (1/K₀). Up to 10 PASEF MS/MS frames were acquired per cycle on ion-mobility-separated precursors, excluding singly charged ions, which were fully resolved in the mobility dimension. Intensity thresholds were set at 1750 counts (minimum) and 14,500 counts (target).

Raw mass spectral data were processed using MetaLab v1.1 (Cheng et al. 2017), employing the embedded FragPipe (v23.0) algorithm (Kong et al. 2017). Searches were conducted against the Chicken Gut MAG database (v1.0.1; 1322 species) (Cheng et al. 2023), supplemented with the UniProt chicken protein database (18,370 entries). MSFragger search parameters included a precursor and fragment mass tolerance of 20 ppm, trypsin enzyme specificity allowing up to two missed cleavages, and a minimum peptide length of seven amino acids. Carbamidomethylation of cysteine was specified as a fixed modification, and methionine oxidation was included as a variable modification. Differential expression analysis of the proteins was performed using DEP/limma, after filtering proteins with filter_missval() (threshold = 0) and identifying significant hits using an adjusted P value < 0.05 and a log₂ fold-change of 1.5. A heatmap of proteins showing significant differences (p<0.05) was plotted using the ggplot2 package.

### RNAseq analysis

The fastaq files of eukaryotic mRNA sequencing data were run through the nf-core/rnaseq (v3.14.0) pipeline using nextflow (v23.10.1) (Ewels et al. 2020). The reads were aligned against the chicken genome (bGalGal1.mat.broiler.GRCg7b) using STAR (v2.7.9a) and gene-level quantification was performed using Salmon (v1.10.1) (https://github.com/COMBINE-lab/salmon) to generate gene counts. Differential expression analysis was performed using DESeq2 (v1.38.3).

## Statistical analysis

Data normality was checked using the Shapiro-Wilk test. The significance of alpha diversity was analysed using the Kruskal-Wallis test, while the Dunn test was performed for multiple comparisons. Ordination analysis of Bray-Curtis distance was visualized using Principal Coordinate analysis. Permutational Multivariate Analysis of Variance (PERMANOVA) with 999 permutations was performed for microbial composition and function using the Adonis2 function from the vegan (v2.6.4) package. Graphs were visualised using ggplot2 (v3.5.0) (Ginestet 2011). The P values were adjusted for the false discovery rate using Benjamini-Hochberg method, and significance was defined as p≤0.05.

## Data availability

The datasets generated and analysed during the current study (fastq files, assemblies, genome bins, and metagenome assembled genomes) are available in the European Nucleotide Archive under the project number PRJEB77488.

## Supporting information

Additional_file_1.docx

## Acknowledgment

The authors acknowledge Ian Hollows from Target Feeds for feed design and formulation. We also acknowledge all of the staff at the National Avian Research Facility for the care of our animals. This work was supported by the BBSRC Institute through Strategic Program Programme Grant funding (BBS/E/RL/230001A). Laura Glendinning is supported by a University of Edinburgh Chancellor’s Fellowship. SG was partially funded by the BBSRC Core Capability Grant BB/CCG2270/1 awarded to the Roslin Institute.

## Contributions

LG, KW, and FK contributed to the study design and provided critical suggestions and feedback. AA, FK, and LG contributed to sample collection. AA and LG contributed to data analysis and interpretation. AA contributed to sample processing, image generation, and manuscript preparation. DK and RK contributed to the metaproteomics data analysis. AA and SG contributed to RNAseq data analysis. All authors read and approved the final manuscript.

## Supplementary Information

### Additional_file_1.docx

Additional file 1: Fig. S1-S5. Fig. S1. Metagenomic (A and C) and metatranscriptomic (B and D) reads classified at the kingdom and phylum levels using Kraken2 across different dietary groups. Only the top 10 most abundant taxa are shown, while the remaining taxa are grouped as other. Reads that couldn’t be assigned any taxonomy are classified as unassigned. CG, Control group; IN1, 1% inulin; IN4, 4% inulin; CE1, 1% ARBOCEL; CE4, 4% ARBOCEL. Fig. S2: Heatmap showing log fold change of significantly differentially abundant caecal microbiota at phylum (A), genus (B), and species (C) levels in pairwise comparisons between different dietary groups. In the pairwise comparisons, the first group serves as the control, while the latter group is considered the treatment group when calculating the log fold change. Significance was declared at p1≤10.05. CG, Control group; IN1, 1% inulin; IN4, 4% inulin; CE1, 1% ARBOCEL; CE4, 4% ARBOCEL. Fig. S3. Heatmap illustrates the log fold change in differentially expressed genes in pairwise comparison of different dietary groups. Significance was declared at p1≤10.05. CG, Control group; IN1, 1% inulin; IN4, 4% inulin; CE1, 1% ARBOCEL; CE4, 4% ARBOCEL. Fig. S4. Principal coordinate analysis (PCoA) of the CAZyme glycoside hydrolases (A) and glycosyltransferases (B) families calculated by Bray-Curtis distance. Each symbol represents an individual sample, with differing symbols and colours indicating the same dietary group. Fig. S5. Heatmap showing the log fold change in differentially expressed proteins in pairwise comparison of different dietary groups. Significance was declared at p ≤10.05. CG, Control group; IN1, 1% inulin; IN4, 4% inulin; CE1, 1% ARBOCEL; CE4, 4% ARBOCEL.

### Additional_file_2.xlsx

Additional file 2: Table S1-S5. Table S1. Total gene counts of CAZyme classes identified in metatranscriptomics data. Table S2. Influence of dietary fibres on the expression of immune-associated genes in chicken caecal epithelium. Table S3. Raw ingredients and nutritional compositions of starter and grower diets across different dietary groups. Table S4. Details of each MAG. Table S5. MAGs showing higher gene counts of CAZymes in IN4 and CE4 groups compared to the CG group.

## References

Abdel-Hamid AM, Attwood MM, Guest JR. 2001. Pyruvate oxidase contributes to the aerobic growth efficiency of Escherichia coli. Microbiology (N Y). 147:1483–1498. doi: 10.1099/00221287-147-6-1483.

Abdelqader A, Al-Fataftah A-R, Daş G. 2013. Effects of dietary Bacillus subtilis and inulin supplementation on performance, eggshell quality, intestinal morphology and microflora composition of laying hens in the late phase of production. Anim Feed Sci Technol. 179:103–111. doi: 10.1016/j.anifeedsci.2012.11.003.

Anand A, Manjula SN, Fuloria NK, Sharma H, Mruthunjaya K. 2025. Inulin as a Prebiotic and Its Effect on Gut Microbiota. In: Inulin for Pharmaceutical Applications. Springer Nature Singapore: Singapore pp. 113–135. doi: 10.1007/978-981-97-9056-2_6.

Anders S, Pyl PT, Huber W. 2015. HTSeq—a Python framework to work with high-throughput sequencing data. bioinformatics. 31:166–169. doi: 10.1093/bioinformatics/btu638.

Aroney STN et al. 2025. CoverM: Read alignment statistics for metagenomics. arXiv preprint arXiv:2501.11217. doi: 10.1093/bioinformatics/btaf147.

Bist RB et al. 2024. Sustainable poultry farming practices: a critical review of current strategies and future prospects. Poult Sci. 103:104295. doi: 10.1016/j.psj.2024.104295.

Chastain CJ et al. 2011. Functional evolution of C4 pyruvate, orthophosphate dikinase. J Exp Bot. 62:3083–3091. doi: 10.1093/jxb/err058.

Chaumeil P-A, Mussig AJ, Hugenholtz P, Parks DH. 2022. GTDB-Tk v2: memory friendly classification with the genome taxonomy database. Bioinformatics. 38:5315–5316. doi: 10.1093/bioinformatics/btac672.

Chen M et al. 2025. Effect of inulin supplementation in maternal fecal microbiota transplantation on the early growth of chicks. Microbiome. 13:98. doi: 10.1186/s40168-025-02084-z.

Chen S. 2023. Ultrafast one-pass FASTQ data preprocessing, quality control, and deduplication using fastp. iMeta. 2:e107. doi: 10.1002/imt2.107.

Cheng K et al. 2017. MetaLab: an automated pipeline for metaproteomic data analysis. Microbiome. 5:157. doi: 10.1186/s40168-017-0375-2.

Cheng K et al. 2023. MetaLab-MAG: A Metaproteomic Data Analysis Platform for Genome-Level Characterization of Microbiomes from the Metagenome-Assembled Genomes Database. J Proteome Res. 22:387–398. doi: 10.1021/acs.jproteome.2c00554.

Chklovski A, Parks DH, Woodcroft BJ, Tyson GW. 2024. Author Correction: CheckM2: a rapid, scalable and accurate tool for assessing microbial genome quality using machine learning. Nat Methods. 21:735. doi: 10.1038/s41592-024-02248-z.

Danecek P et al. 2021. Twelve years of SAMtools and BCFtools. Gigascience. 10:giab008. doi: 10.1093/gigascience/giab008.

Evans JT, Denef VJ. 2020. To Dereplicate or Not To Dereplicate? mSphere. 5. doi: 10.1128/mSphere.00971-19.

Ewels PA et al. 2020. The nf-core framework for community-curated bioinformatics pipelines. Nat Biotechnol. 38:276–278. doi: 10.1038/s41587-020-0439-x.

FAO. 2023. Gateway to poultry production and products.

FAO. 2021. The State of Food Security and Nutrition in the World 2021. FAO, IFAD, UNICEF, WFP and WHO doi: 10.4060/cb4474en.

Farkas V et al. 2025. Even Low Amounts of Amorphous Lignocellulose Affect Some Upper Gut Parameters, but They Do Not Modify Ileal Microbiota in Young Broiler Chickens. Animals. 15:851. doi: 10.3390/ani15060851.

Farris J, Morgan S, Beckman J. 2024. Evaluating the effects of nontariff measures on poultry trade. doi: 10.32747/2024.8453400.ers.

Feng Y et al. 2025. Advances in understanding dietary fiber: Classification, structural characterization, modification, and gut microbiome interactions. Compr Rev Food Sci Food Saf. 24. doi: 10.1111/1541-4337.70092.

Gemayel K, Lomsadze A, Borodovsky M. 2022. MetaGeneMark-2: improved gene prediction in metagenomes. BioRxiv. 2022–2027. doi: 10.1101/2022.07.25.500264.

Ginestet C. 2011. ggplot2: Elegant Graphics for Data Analysis. J R Stat Soc Ser A Stat Soc. 174:245–246. doi: 10.1111/j.1467-985X.2010.00676_9.x.

Govoni C et al. 2021. Global assessment of natural resources for chicken production. Adv Water Resour. 154:103987. doi: 10.1016/j.advwatres.2021.103987.

Hou L, Sun B, Yang Y. 2020. Effects of Added Dietary Fiber and Rearing System on the Gut Microbial Diversity and Gut Health of Chickens. Animals. 10:107. doi: 10.3390/ani10010107.

Hyatt D et al. 2010. Prodigal: prokaryotic gene recognition and translation initiation site identification. BMC Bioinformatics. 11:1–11. doi: 10.1186/1471-2105-11-119.

Jamroz D, Jakobsen K, Knudsen KEB, Wiliczkiewicz A, Orda J. 2002. Digestibility and energy value of non-starch polysaccharides in young chickens, ducks and geese, fed diets containing high amounts of barley. Comp Biochem Physiol A Mol Integr Physiol. 131:657–668. doi:/10.1016/S1095-6433(01)00517-7.

Jha R, Mishra P. 2021. Dietary fiber in poultry nutrition and their effects on nutrient utilization, performance, gut health, and on the environment: a review. J Anim Sci Biotechnol. 12:51. doi: 10.1186/s40104-021-00576-0.

Jian Z et al. 2025. Species and functional composition of cecal microbiota and resistance gene diversity in different Yunnan native chicken breeds: A metagenomic analysis. Poult Sci. 104:105138. doi: 10.1016/j.psj.2025.105138.

Kaenying W et al. 2023. Structural and mutational analysis of glycoside hydrolase family 1 Br2 β-glucosidase derived from bovine rumen metagenome. Heliyon. 9:e21923. doi: 10.1016/j.heliyon.2023.e21923.

Kang DD et al. 2019. MetaBAT 2: an adaptive binning algorithm for robust and efficient genome reconstruction from metagenome assemblies. PeerJ. 7:e7359. doi: 10.7717/peerj.7359.

Kong AT, Leprevost F V, Avtonomov DM, Mellacheruvu D, Nesvizhskii AI. 2017. MSFragger: ultrafast and comprehensive peptide identification in mass spectrometry–based proteomics. Nat Methods. 14:513–520. doi: 10.1038/nmeth.4256.

Kopylova E, Noé L, Touzet H. 2012. SortMeRNA: fast and accurate filtering of ribosomal RNAs in metatranscriptomic data. Bioinformatics. 28:3211–3217. doi: 10.1093/bioinformatics/bts611.

Li D, Liu C-M, Luo R, Sadakane K, Lam T-W. 2015. MEGAHIT: an ultra-fast single-node solution for large and complex metagenomics assembly via succinct de Bruijn graph. Bioinformatics. 31:1674–1676. doi: 10.1093/bioinformatics/btv033.

Li H. 2013. Aligning sequence reads, clone sequences and assembly contigs with BWA-MEM. arXiv preprint arXiv:1303.3997. doi: 10.48550/arXiv.1303.3997.

Li H et al. 2009. The sequence alignment/map format and SAMtools. Bioinformatics. 25:2078–2079. doi: 10.1093/bioinformatics/btp352.

Lin H, Das Peddada S. 2020. Analysis of compositions of microbiomes with bias correction. Nat Commun. 11:3514. doi: 10.1038/s41467-020-17041-7.

Liu HY, Hou R, Yang GQ, Zhao F, Dong WG. 2018. In vitro effects of inulin and soya bean oligosaccharide on skatole production and the intestinal microbiota in broilers. J Anim Physiol Anim Nutr (Berl). 102:706–716. doi: 10.1111/jpn.12830.

Lu J, Breitwieser FP, Thielen P, Salzberg SL. 2017. Bracken: estimating species abundance in metagenomics data. PeerJ Comput Sci. 3:e104. doi: 10.7717/peerj-cs.104.

Makki K, Deehan EC, Walter J, Bäckhed F. 2018. The Impact of Dietary Fiber on Gut Microbiota in Host Health and Disease. Cell Host Microbe. 23:705–715. doi: 10.1016/j.chom.2018.05.012.

Marc RA et al. 2024. Dietary Fibers and Their Importance in the Diet. IntechOpen. doi: 10.5772/intechopen.115461.

McMurdie PJ, Holmes S. 2013. phyloseq: An R Package for Reproducible Interactive Analysis and Graphics of Microbiome Census Data. PLoS One. 8:e61217-. doi: 10.1371/journal.pone.0061217.

Mirzaie S, Zaghari M, Aminzadeh S, Shivazad M. 2012. The effects of non-starch polysaccharides content of wheat and xylanase supplementation on the intestinal amylase, amino peptidase and lipase activities, ileal viscosity and fat digestibility in layer diet.

Nguyen HT, Bedford MR, Morgan NK. 2021. Importance of considering non-starch polysaccharide content of poultry diets. Worlds Poult Sci J. 77:619–637. doi: 10.1080/00439339.2021.1921669.

Nguyen HT, Bedford MR, Wu S-B, Morgan NK. 2022. Dietary soluble non-starch polysaccharide level influences performance, nutrient utilisation and disappearance of non-starch polysaccharides in broiler chickens. Animals. 12:547. doi: 10.3390/ani12050547.

Nicholls C, Li H, Liu J. 2012. GAPDH: A common enzyme with uncommon functions. Clin Exp Pharmacol Physiol. 39:674–679. doi: 10.1111/j.1440-1681.2011.05599.x.

Novoa Rama E et al. 2023. Characterizing the gut microbiome of broilers raised under conventional and no antibiotics ever practices. Poult Sci. 102:102832. doi: 10.1016/j.psj.2023.102832.

Ocejo M, Oporto B, Hurtado A. 2019. 16S rRNA amplicon sequencing characterization of caecal microbiome composition of broilers and free-range slow-growing chickens throughout their productive lifespan. Sci Rep. 9:2506. doi: 10.1038/s41598-019-39323-x.

OECD/FAO. 2018. OECD-FAO Agricultural Outlook 2018-2027. OECD doi: 10.1787/agr_outlook-2018-en.

OECD/FAO. 2024. OECD-FAO Agricultural Outlook 2024-2033. Paris/FAO, Rome doi: 10.1787/4c5d2cfb-en.

Pérez-Jiménez J. 2024. Dietary fiber: Still alive. Food Chem. 439:138076. doi: 10.1016/j.foodchem.2023.138076.

Pouyez J et al. 2012. First crystal structure of an endo-inulinase, INU2, from Aspergillus ficuum: Discovery of an extra-pocket in the catalytic domain responsible for its endo-activity. Biochimie. 94:2423–2430. doi: 10.1016/j.biochi.2012.06.020.

Qiu M et al. 2022. Research Note: The gut microbiota varies with dietary fiber levels in broilers. Poult Sci. 101:101922. doi: 10.1016/j.psj.2022.101922.

Shen W, Le S, Li Y, Hu F. 2016. SeqKit: a cross-platform and ultrafast toolkit for FASTA/Q file manipulation. PLoS One. 11:e0163962. doi: 10.1371/journal.pone.0163962.

Shterzer N et al. 2023. Vertical transmission of gut bacteria in commercial chickens is limited. Anim Microbiome. 5:50. doi: 10.1186/s42523-023-00272-6.

Song J et al. 2020. Dietary Inulin Supplementation Modulates Short-Chain Fatty Acid Levels and Cecum Microbiota Composition and Function in Chickens Infected With Salmonella. Front Microbiol. 11. doi: 10.3389/fmicb.2020.584380.

de Sousa LS et al. 2025. Cecal microbial composition and serum concentration of short-chain fatty acids in laying hens fed different fiber sources. Brazilian Journal of Microbiology. 1–14. doi: 10.1007/s42770-024-01606-5.

Suzek BE et al. 2015. UniRef clusters: a comprehensive and scalable alternative for improving sequence similarity searches. Bioinformatics. 31:926–932. doi: 10.1093/bioinformatics/btu739.

Wood DE, Lu J, Langmead B. 2019. Improved metagenomic analysis with Kraken 2. Genome Biol. 20:1–13. doi: 10.1186/s13059-019-1891-0.

Xia Y et al. 2021. Dietary inulin supplementation modulates the composition and activities of carbohydrate-metabolizing organisms in the cecal microbiota of broiler chickens. PLoS One. 16:e0258663. doi: 10.1371/journal.pone.0258663.

Yasmeen R, Ahmad F. 2025. Microbial fermented agricultural waste-based broiler feed: a sustainable alternative to conventional feed. Worlds Poult Sci J. 81:271–287. doi: 10.1080/00439339.2024.2443222.

Zeng C, Zeng X, Xia S, Ye G. 2024. Caproicibacterium argilliputei sp. nov., a novel caproic acid producing anaerobic bacterium isolated from pit clay. Int J Syst Evol Microbiol. 74. doi: 10.1099/ijsem.0.006246.

Zheng J et al. 2023. dbCAN3: automated carbohydrate-active enzyme and substrate annotation. Nucleic Acids Res. 51:W115–W121. doi: 10.1093/nar/gkad328.

