## Additional_file_1.docx for "Integrative multi-omics analysis of dietary fibre-induced modulations in the composition and function of chicken caecal microbiota"

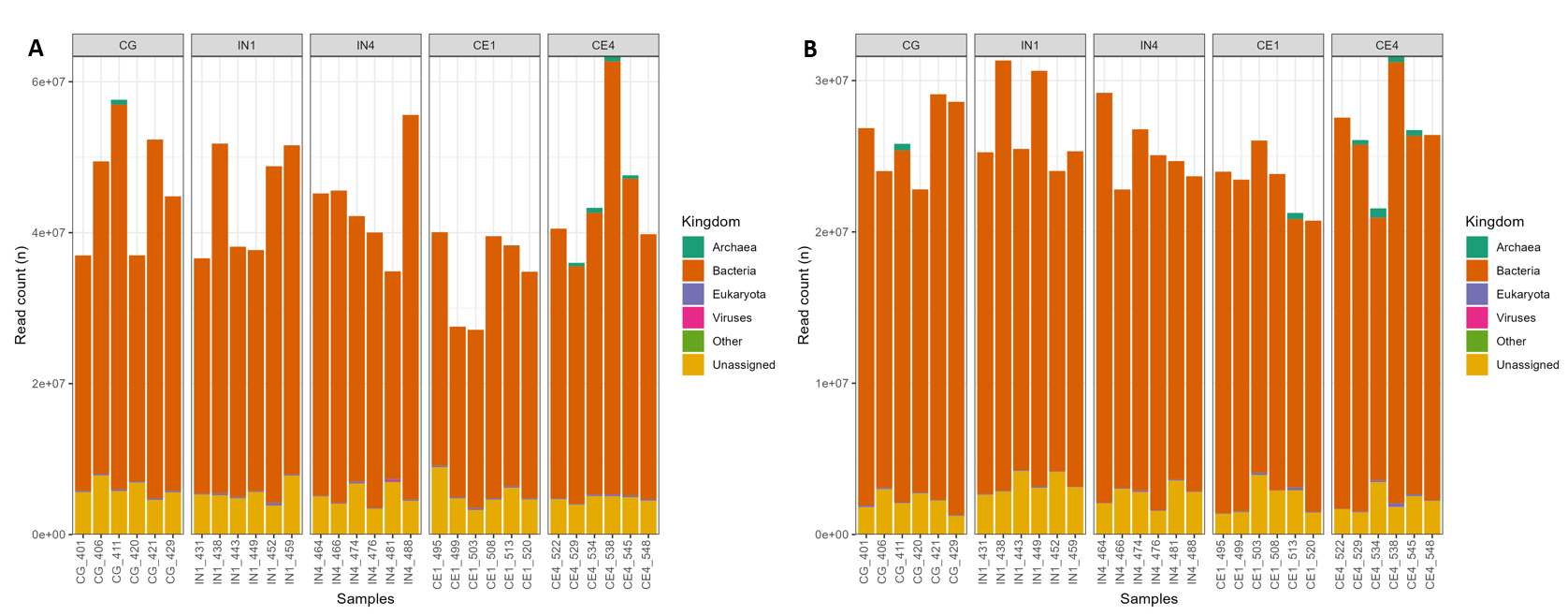

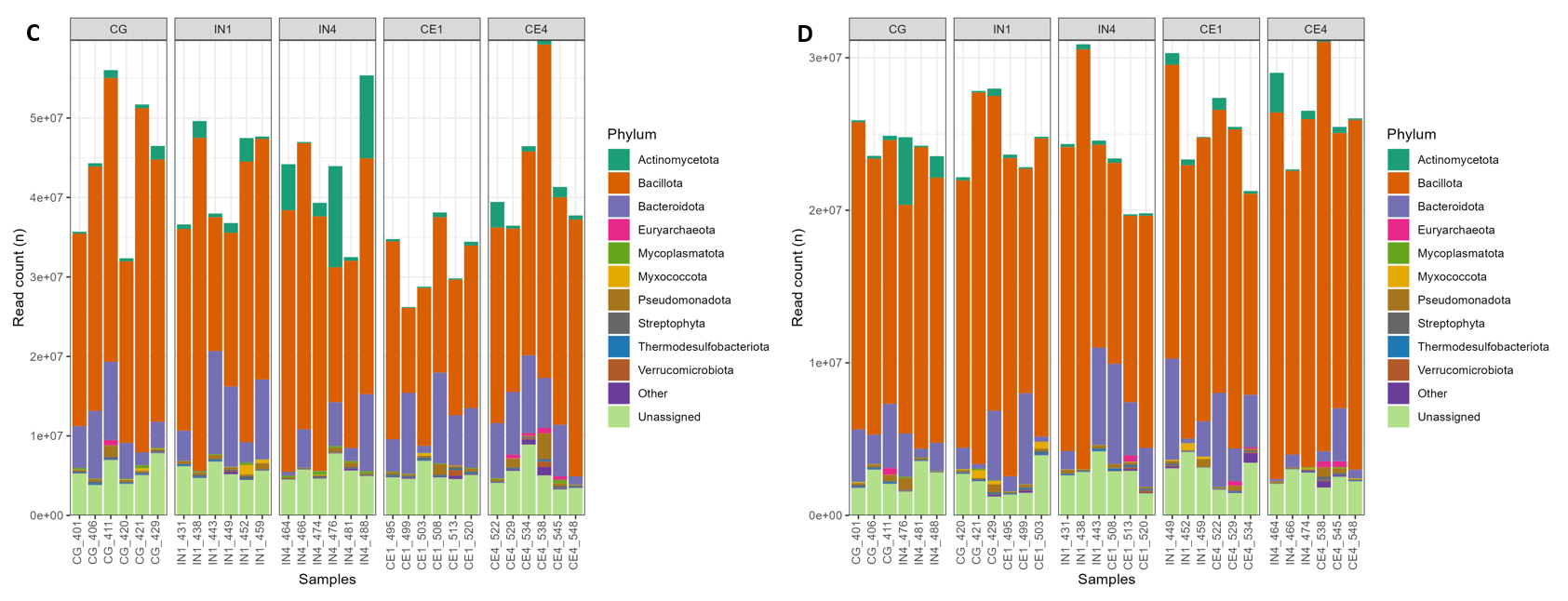

Fig. S1. Metagenomic (A and C) and metatranscriptomic (B and D) reads classified at the kingdom and phylum levels using Kraken2 across different dietary groups. Only the top 10 most abundant taxa are shown, while the remaining taxa are grouped as other. Reads that couldn’t be assigned any taxonomy are classified as unassigned. CG, Control group; IN1, 1% inulin; IN4, 4% inulin; CE1, 1% ARBOCEL; CE4, 4% ARBOCEL.

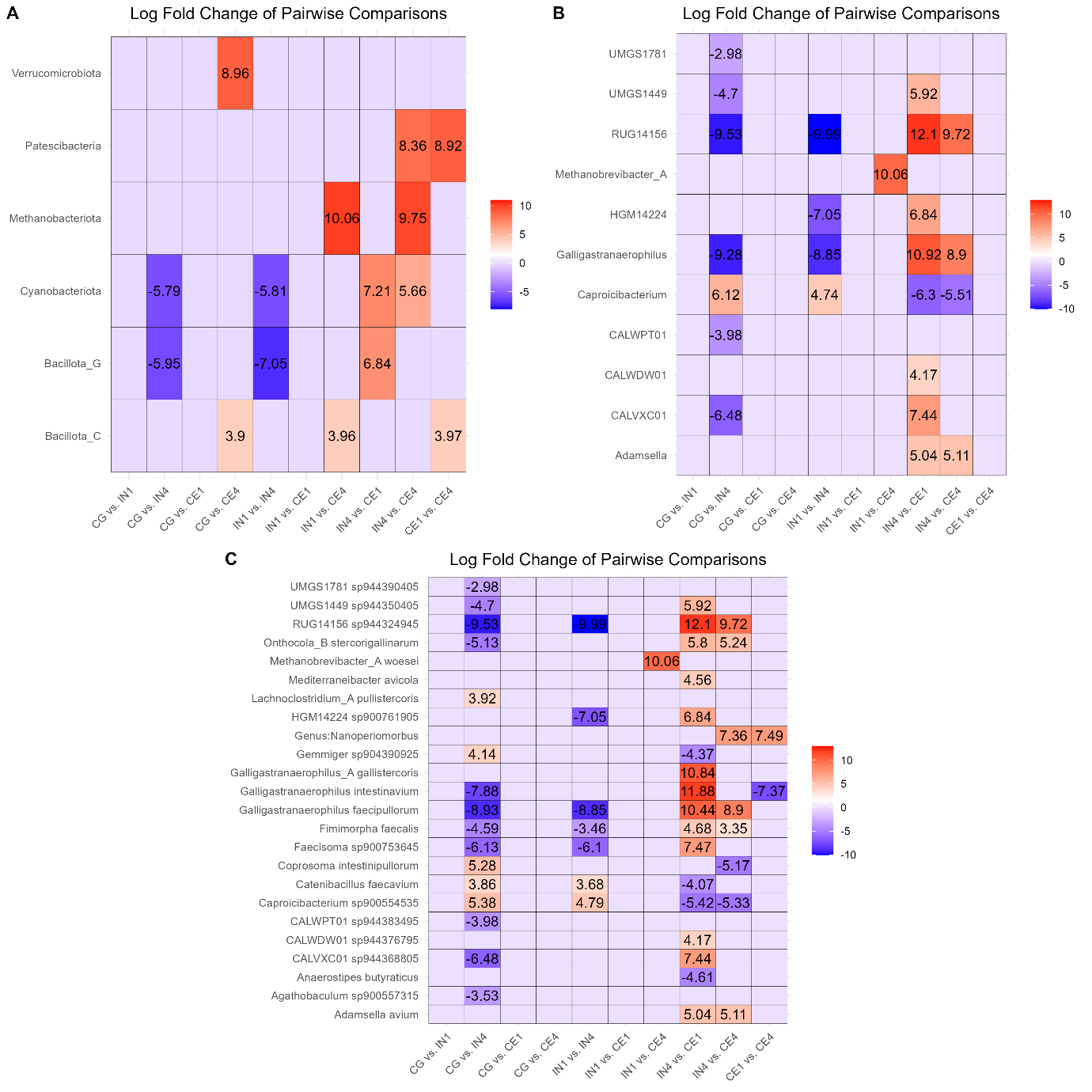

Fig. S2: Heatmap showing log fold change of significantly differentially abundant caecal microbiota at phylum (A), genus (B), and species (C) levels in pairwise comparisons between different dietary groups. In the pairwise comparisons, the first group serves as the control, while the latter group is considered the treatment group when calculating the log fold change. Significance was declared at p ≤ 0.05. CG, Control group; IN1, 1% inulin; IN4, 4% inulin; CE1, 1% ARBOCEL; CE4, 4% ARBOCEL.

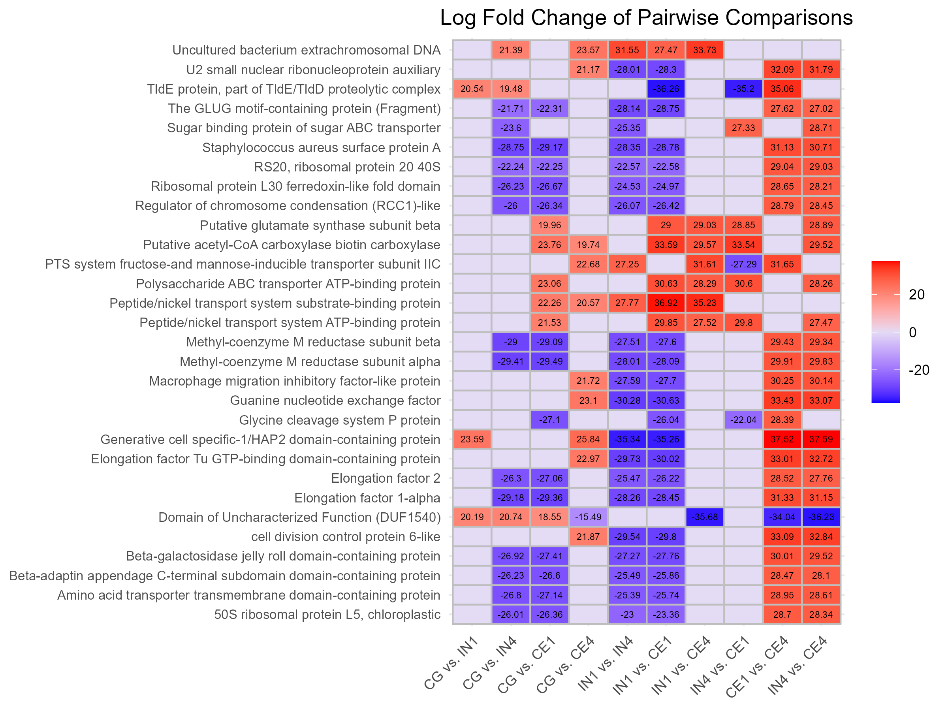

Fig. S3. Heatmap illustrates the log fold change in differentially expressed genes in pairwise comparison of different dietary groups. Significance was declared at p ≤ 0.05. CG, Control group; IN1, 1% inulin; IN4, 4% inulin; CE1, 1% ARBOCEL; CE4, 4% ARBOCEL.

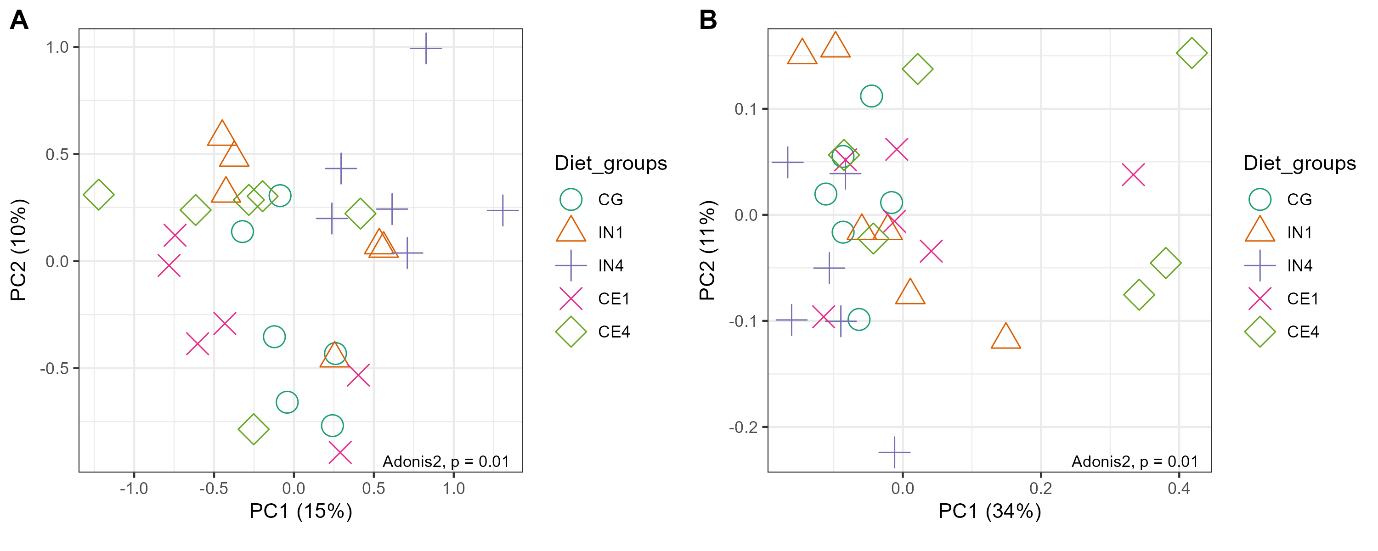

Fig. S4. Principal coordinate analysis (PCoA) of the CAZyme glycoside hydrolases (A) and glycosyltransferases (B) families calculated by Bray-Curtis distance. Each symbol represents an individual sample, with differing symbols and colours indicating the same dietary group. CG, Control group; IN1, 1% inulin; IN4, 4% inulin; CE1, 1% ARBOCEL; CE4, 4% ARBOCEL.

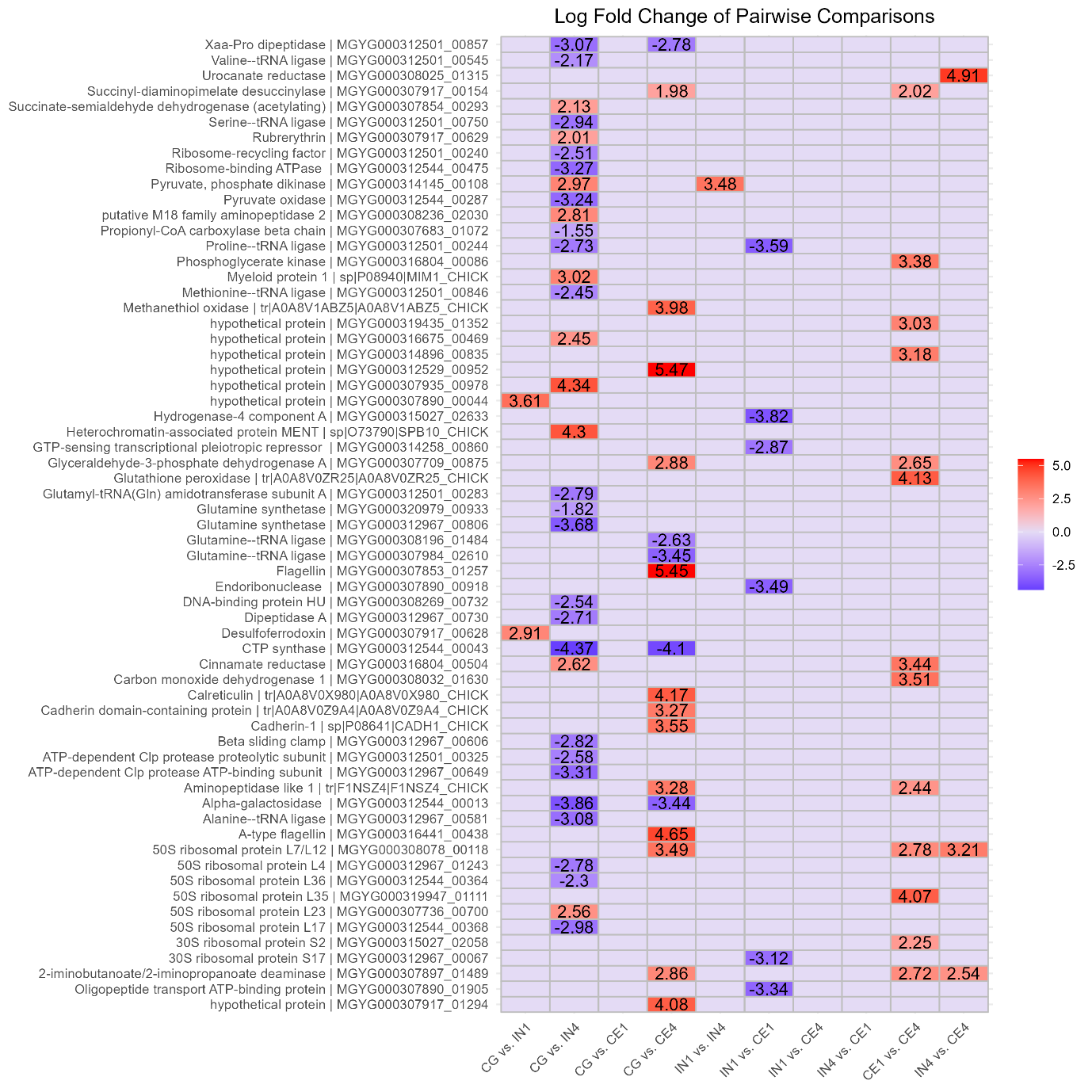

Fig. S7. Heatmap showing the log fold change in differentially expressed proteins in pairwise comparison of different dietary groups. Significance was declared at p ≤ 0.05. CG, Control group; IN1, 1% inulin; IN4, 4% inulin; CE1, 1% ARBOCEL; CE4, 4% ARBOCEL.
